# Photophysical studies at cryogenic temperature reveal a novel photoswitching mechanism of rsEGFP2

**DOI:** 10.1101/2022.08.22.504779

**Authors:** Angela M. R. Mantovanelli, Oleksandr Glushonkov, Virgile Adam, Jip Wulffele, Daniel Thédié, Martin Byrdin, Ingo Gregor, Oleksii Nevskyi, Jörg Enderlein, Dominique Bourgeois

## Abstract

Single-molecule-localization-microscopy (SMLM) at cryogenic temperature opens new avenues to investigate intact biological samples at the nanoscale and perform cryo-correlative studies. Genetically encoded fluorescent proteins (FPs) are markers of choice for cryo-SMLM, but their reduced conformational flexibility below the glass transition temperature hampers efficient photoswitching at low temperature. We investigated cryo-switching of rsEGFP2, one of the most efficient reversibly switchable fluorescent protein at ambient temperature due to facile *cis-trans* isomerization of the chromophore. UV-visible microspectrophotometry and X-ray crystallography revealed a completely different switching mechanism at ∼110 K. At this cryogenic temperature, on-off photoswitching involves the formation of 2 dark states with blue shifted absorption relative to that of the *trans* protonated chromophore populated at ambient temperature. Only one of these dark states can be switched back to the fluorescent state by 405 nm light, while both of them are sensitive to UV light at 355 nm. The rsEGFP2 photoswitching mechanism discovered in this work adds to the panoply of known switching mechanisms in fluorescent proteins. It suggests that employing 355 nm light in cryo-SMLM experiments using rsEGFP2 or possibly other FPs could improve the achievable effective labeling efficiency in this technique.

**Table of Contents artwork:** 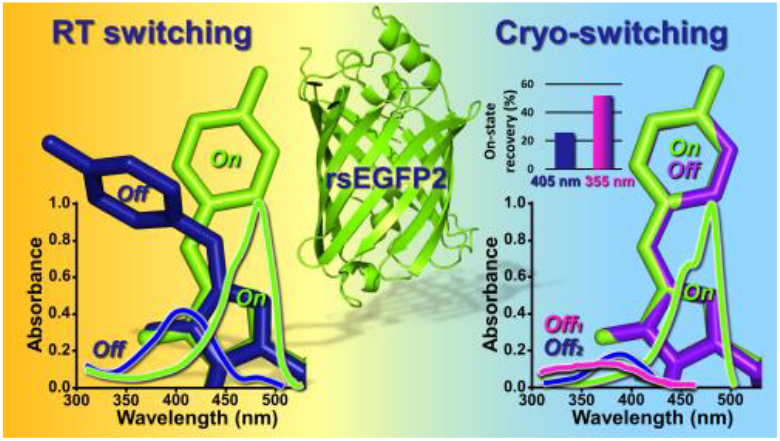

## INTRODUCTION

Super-resolution fluorescence microscopy has revolutionized our ability to investigate life at the nanoscale ^1^. Yet, to prevent motion artifacts and facilitate labeling, many nanoscopy studies are still based on chemically fixed cells. Chemical fixation of membrane proteins remains challenging, and a number of artifacts may result from fixation that become even more detrimental as the quest for high-resolution increases ^2–4^. One strategy to better preserve the fine morphological details of biological samples is to image flash-frozen cells at cryogenic temperature (CT) ^5–14^. Furthermore, performing super-resolution microscopy at CT opens the door to cryo-correlative (CLEM) studies with cryo-electron microscopy ^6,10,15^. In addition, CT-fluorescence imaging offers additional benefits such as improved quantum yield and reduced photobleaching of the fluorescence markers, as well as narrowing of fluorescence emission bands, potentially facilitating multicolor data acquisition schemes ^6,11,16–18^. Yet, although impressive progresses have been achieved recently ^14,19,20^, cryo-nanoscopy faces significant challenges. One of those concerns the development of user-friendly instrumentation compatible with high numerical-aperture objectives and long data acquisition time with low sample drift ^11,21–24^. A more fundamental issue concerns the development of fluorescent markers with efficient photoswitching properties at CT ^7,13,17,25–27^.

Fluorescent proteins (FPs) are arguably the most appropriate markers for super-resolution microscopy at CT, as they are genetically encoded and thus do not require fixation or permeabilization for efficient labeling. In contrast, many organic dyes do not cross membranes naturally. Furthermore, organic dyes typically used in single-molecule localization techniques such as stochastic optical reconstruction microscopy ((d)STORM) or point accumulation in nanoscale topography (PAINT) cannot be used at CT because their efficient blinking relies on the diffusion of buffer molecules or of the fluorophores themselves, which is arrested in a frozen solvent. A number of fluorescent proteins have been tested for their ability to switch at CT ^7–9,11,13,14,25,27^. While phototransformable FPs ^28,29^ have been mostly investigated ^7,8,11,25–27^, other more standard FPs have also been shown to undergo cryo-switching ^8,9,14,26^. Conflicting results have sometimes been reported, notably concerning reversibly switchable FPs (RSFPs). For example Dronpa has been shown to not switch at CT, due to restricted structural dynamics ^7,30^, to cryo-switch moderately ^26^ or quite efficiently ^8,13^. While the positively photoswitchable FP Padron was reported to maintain *trans* to *cis* isomerization at 100 K ^27^, negative photoswitching at 77 K was also observed ^13^. Overall, mechanistic knowledge about the photoswitching mechanisms adopted by FPs at CT remains scarce, although some hypotheses have been put forward such as the possible involvement of the triplet state in the case of mEmerald ^14^ or of Kolbe-driven photo-decarboxylation of the conserved Glu222 (GFP amino-acid numbering) in PA-GFP or PA-mKate ^25,31^. In this work, employing a combination of UV-visible microspectrophotometry, X-ray crystallography and single-molecule studies, we focused on the cryo-switching mechanism of rsEGFP_2_, a well-known RSFP ^32^ for which extended knowledge has been gathered in the case of switching at room temperature (RT) ^33–37^. We show that switching of rsEGFP2 at ∼110 K proceeds via a completely different pathway than at RT, which does not involve chromophore isomerization but instead populates 2 dark states in the *cis* conformation of the chromophore. Based on ensemble data and advanced single-molecule simulations, we show that the use of 355 nm laser light is expected to enhance the effective rsEGFP_2_ labeling efficiency in cryo-PALM experiments.

## RESULTS

Stimulated by our recent structural investigations of rsEGFP2 at RT ^33–36^, and by the fact that this RSFP was recently shown to maintain efficient switching at 77 K ^13^, we were interested to know whether cryo-switching proceeds by the same *cis trans* isomerization mechanism as observed at RT. In view of the very efficient switching of rsEGFP2 observed at 300 K, we initially assumed that sufficient conformational flexibility could be maintained at CT to enable isomerization. UV-visible microspectrophotometry experiments using a dedicated instrument ^38^ were first performed on flash-cooled samples of purified rsEGFP2 mixed with glycerol which were held in micro capillaries (Supplementary Methods).

In comparison with RT switching, a considerably reduced rate of switching upon illumination with 488 nm laser light was observed at ∼110 K (Supplementary Fig. 1, Supplementary Fig. 2). Furthermore, in contrast with the off-state reached at RT (absorption peak maximum at 411 nm) (Fig. 1A), the absorption peak for the cryo-switched off-state (Fig. 1C) was largely blue shifted (peak maximum at 385 nm) and more structured. The blueshift was not an effect of the temperature at which spectra were recorded, as the absorption of a sample switched at RT followed by flash cooling was only blue shifted to a minor extent (peak maximum at 406 nm) (Fig. 1E). Whereas back switching by typical 405 nm laser light was nearly complete at RT (Fig. 1B), only partial recovery was obtained at CT after extensive illumination (Fig. 1D, Supplementary Fig. 1). A significant fraction of rsEGFP2 molecules appeared to be trapped in the off-state, and a minor red-shifted absorbance peak was also observed to grow at 520 nm. Interestingly, if rsEGFP2 was off-switched at RT and then flash cooled to CT (Fig. 1E), little back switching could be observed by 405 nm illumination, populating mostly the red shifted absorbance peak (Fig. 1F). This suggests that below the glass transition temperature the protonated *trans* chromophore (the off-switched state at RT) is unable to efficiently undergo *trans* to *cis* back-isomerization followed by deprotonation to the canonical anionic fluorescent state absorbing at 479 nm. Taken together, these data suggest that the off-state reached upon 488 nm illumination at CT differs from that populated at RT.

**Figure 1:**
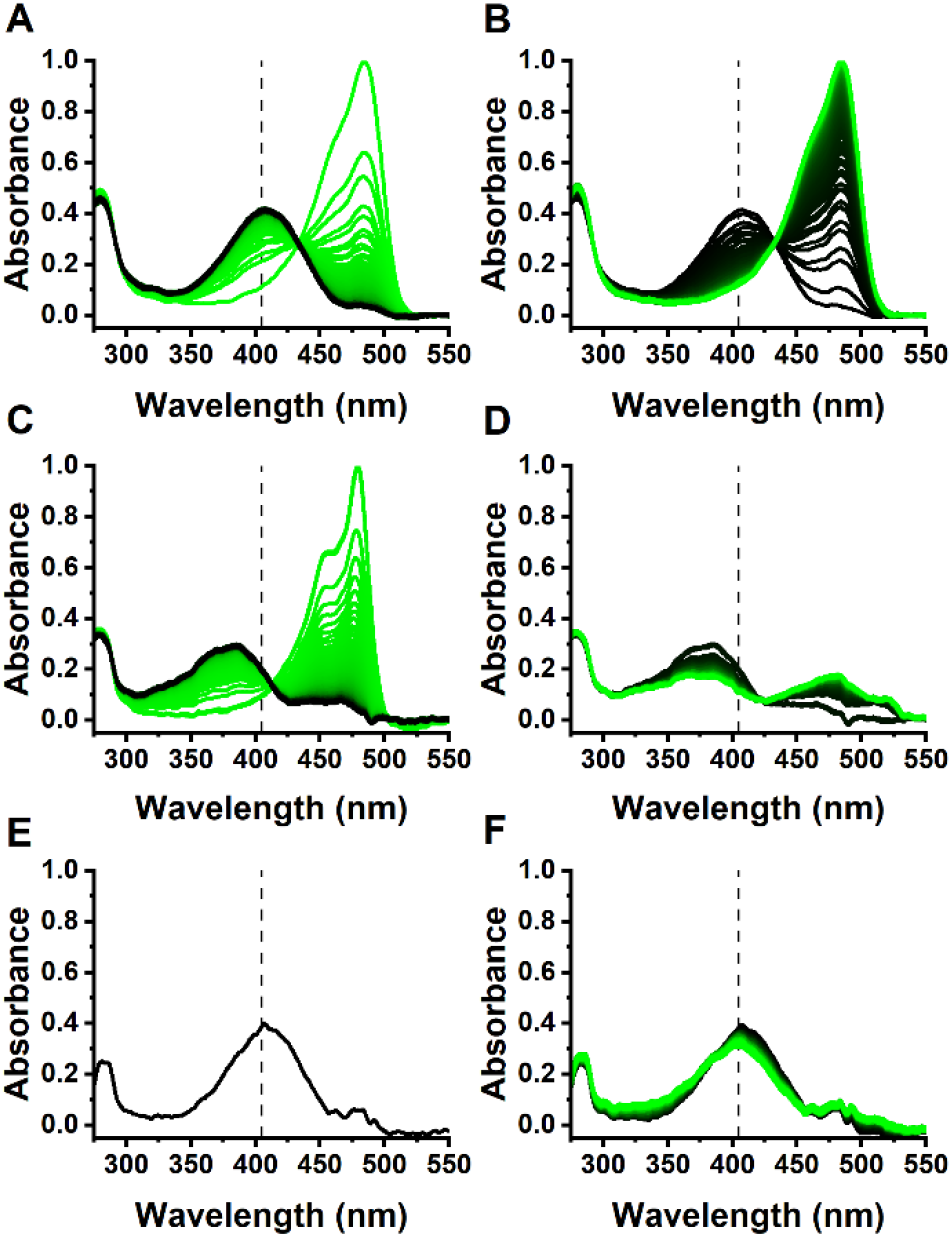
Photoswitching of rsEGFP_2_ monitored by absorption microspectrophotometry. (A) on-to-off switching at RT with 488 nm laser light (80 W/cm^2^). (B) off-to-on switching at RT with 405 nm laser light (4 W/cm^2^). (C) on-to-off switching at 110 K with 488 nm laser light (5.8 kW/cm2). (D) off-to-on swiching at CT with 405 nm laser light (0.7 kW/cm^2^). (E) onto-off switching at RT followed by flash cooling. (F) on-to-off switching at RT followed by flash cooling and illumination with 405 nm laser light (0.1 kW/cm^2^). Spectral series evolve from green to black during off-switching (A, C) or from black to green during on-switching (B, D, F). Absorbance spectra in (A, B) and (C, D) were normalized at the anionic chromophore peak of the first spectrum of each series in A and C, respectively. Spectra in E and F were normalized to match the height of the protonated peak of the first spectrum in A. Representative spectral series are shown from n ≥ 3 measurements. Dashed vertical lines are positioned at 405 nm to guide the eye.

**Figure 2:**
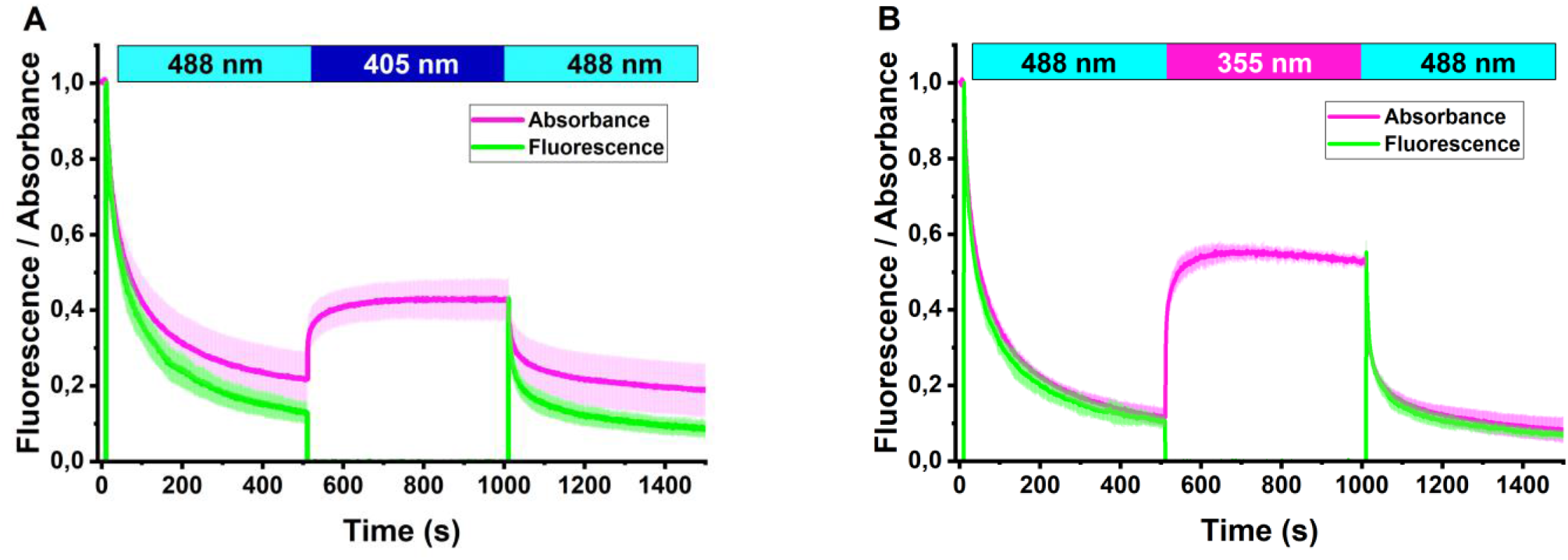
On-state recovery of switched off rsEGFP2 at cryogenic temperature. Absorbance and fluorescence levels are calculated by integration of the absorption and fluorescence emission spectra in 470-500 nm and 495-630 nm spectral ranges, respectively. (A) recovery by 405 nm light. (B) recovery by 355 nm light. Absorbance (magenta) and fluorescence (green) switching kinetics are measured at 110 K by alternate illumination at 488 nm (0.4 kW/cm^2^) and either 405 nm (0.2 kW/cm^2^) or 355 nm (0.03 kW/cm^2^), as indicated in the upper bars. Absorbance and fluorescence are normalized to 1 at start of acquisition. Fluorescence is only measured in the presence of 488 nm light. The mean ± s.d. of n = 3 measurements is shown.

Titration experiments as a function of the employed laser power and fitting of the observed cryo-switching rates indicate that off-switching by 488 nm light as well as on-switching by 405 nm light are (at least) biphasic (Supplementary Fig. 3). The multiphasic nature of photoswitching curves in rsEGFP_2_ has been observed at RT ^36^ and could result from heterogeneous FP populations. It also follows from the fact that at CT the protein molecules cannot tumble in the vitreous solvent and thus switch at different rates depending on their dipole orientation. The fitted rates varied linearly as a function of the applied laser power, suggesting that both off-switching and on-switching at 110 K proceed via single-photon absorption mechanisms (Supplementary Fig. 3).

**Figure 3:**
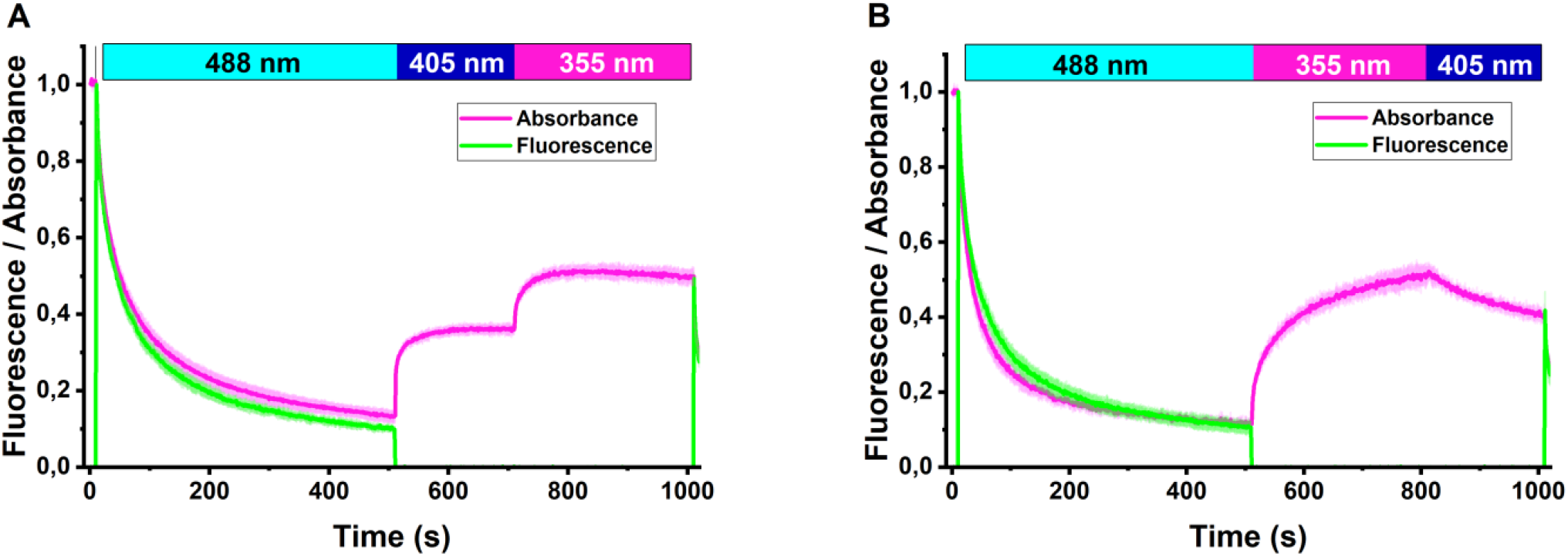
Recovery of rsEGFP2 on state by subsequent illumination with 405 nm and 355 nm light. Absorbance and fluorescence levels are calculated by integration of the absorption and fluorescence emission spectra in 470-500 nm and 495-630 nm spectral ranges, respectively. (A) recovery by 405 nm light followed by 355 nm light. (B) recovery by 355 nm light followed by 405 nm light. Absorbance (magenta) and fluorescence (green) evolutions are measured at 110 K by illumination at 488 nm (0.4 kW/cm^2^), 405 nm (0.3 kW/cm^2^) and 355 nm (0.025 kW/cm^2^), according to the schemes indicated in the upper bars. Absorbance and fluorescence are normalized to 1 at start of acquisition. Fluorescence was only measured in the presence of 488 nm light. The mean ± s.d. of n = 3 measurements is shown.

In line with the absorbance data, off-switching at CT of rsEGFP_2_ by 488 nm light and subsequent on-switching by 405 nm light only allowed the recovery of ∼25 % of the fluorescence (Fig. 2A). Such a low recovery level is problematic for cryo-PALM applications, as only a minor fraction of the fluorescently labeled biological targets would then be detectable, giving rise to a low effective labeling efficiency. In comparison, in standard RT PALM using greento-red photoconvertible FPs, typically 60 to 70% of the labels can be imaged under favorable illumination conditions ^39,40^.

Photobleaching during off-switching by 488 nm light or on switching by 405 nm light at CT can be invoked to explain the low recovery level to the fluorescent state. In particular, increasing the 405 nm light power density resulted in faster recovery that was nevertheless followed by a progressive decay of the on-state absorbance (Supplementary Fig. 4A), suggesting a balance between back switching and photobleaching mechanisms. Yet, the fraction of recovered on-state absorbance was independent of the applied 488 nm or 405 nm power density, suggesting that nonlinear bleaching mechanisms are not predominant (Supplementary Fig. 4B).

**Figure 4:**
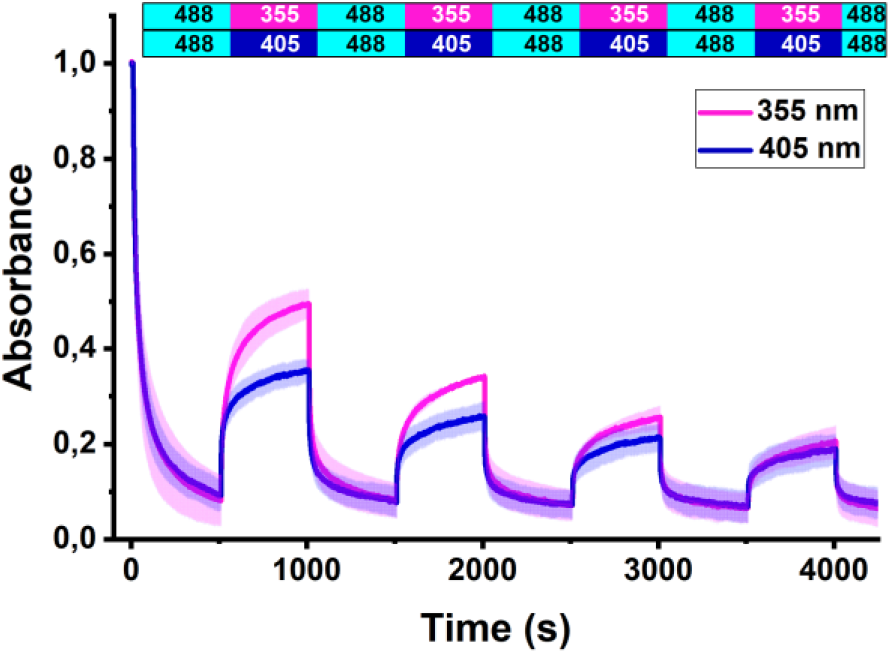
Absorbance photofatigue switching curves of rsEGFP2 at CT. The absorbance signal was calculated by integration of the absorption spectra in the 470-500 nm spectral range. rsEGFP_2_ was switched back and forth with 488 nm (1.0 kW/cm^2^) and either 405 nm (0.2 kW/cm^2^, blue) or 355 nm (0.01 kW/cm2, magenta) laser light, according to the illumination schemes shown in the upper bars. The mean ± s.d. of n = 3 measurements is shown.

In addition to photobleaching, two other mechanisms may contribute to the limited recovery of the fluorescent on state. The first mechanism is similar to that limiting the photoswitching contrast in e.g. RESOLFT experiments at RT ^36^ and involves residual off-switching by 405 nm light. Calculations that assume a wavelength independent off-switching quantum yield (Supplementary Methods) suggest that the ratio of on-switching and off-switching rates by the 405 nm light amounts to ∼900. Therefore, off-switching by 405 nm light is not expected to contribute to the limited recovery level.

The second mechanism involves dark state trapping and would be in line with the residual off-state absorption observed in the spectra of Fig. 1D. In fact, although 405 nm light is nearly centered on the absorption band of the off-switched rsEGFP2 chromophore at RT, it sits on the red edge of the CT off-switched absorption band, possibly limiting on-state recovery. Thus, we replaced the 405 nm laser by a 355 nm laser. This wavelength sits on the blue edge of the absorption band, and the higher energy photons may thus interact more efficiently with the CT off-switched chromophore. 355 nm light effectively enhanced the rsEGFP_2_ recovery level to ∼50 % (Fig. 2B), in line with near-complete disappearance of the off-state absorption band after illumination (Supplementary Fig. 5).

**Figure 5.**
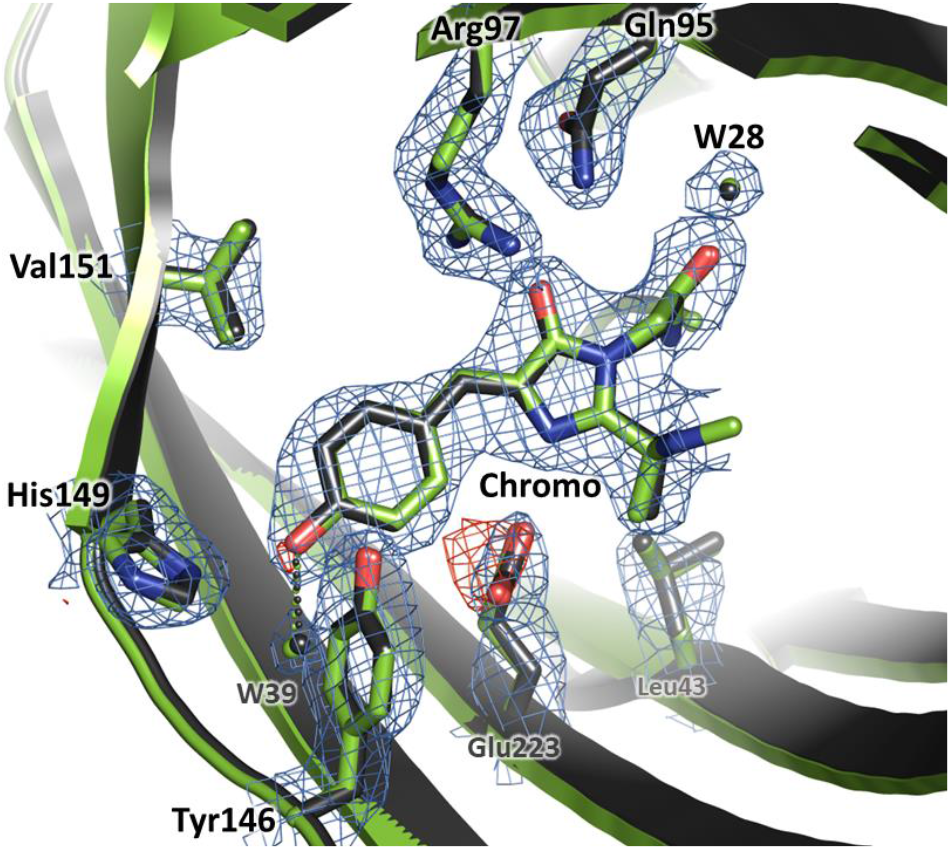
Crystallographic views of rsEGFP2 switching at CT. Refined models of the chromophore and surrounding residues of rsEGFP2 are shown in the *cis* on state (green carbons and water molecules) and in the cryo-switched off state (dark grey carbons and water molecules, PDB code 8AHA), switched with 488 nm laser light for 700 s (0.05 kW/cm2). The 2F_obs_-F_calc_ electron density map contoured at 1σ(blue) and the F_obs_-F_calc_ difference electron density maps contoured at ±3 σ(red: negative, green: positive) of the cryo-switched off state are shown. W: water molecules

To further confirm the different on-switching efficiencies of 405 nm and 355 nm light, we sequentially applied both lasers (Fig. 3). Application of 355 nm light after 405 nm light increased the recovery level to ∼45 %, close to the level observed with 355 nm light only (Fig. 3A). Application of 405 nm light after 355 nm light substantially decreased the recovery level (Fig. 3B). Those data suggest that 355 nm light is able to pump back to the on-state a fraction of rsEGFP_2_ molecules residing in an off-state that does not respond to 405 nm light (*Off*_*1*_), while another fraction of molecules appears to reside in a second off-state (*Off*_*2*_) sensitive to both 405 nm and 355 nm light. Upon off-switching at CT by 488 nm light, *Off*_*1*_ and *Off*_*2*_ are populated and do not exchange significantly. Application of 405 nm light after 355 nm light repopulates *Off*_*1*_ due to residual off-switching at this wavelength while the *Off*_*2*_ steady state level is maintained. This mechanism is clearly visible upon monitoring the absorbance at 320 nm (Supplementary Fig. 6). The observation of *Off*_*1*_ and *Off*_*2*_ is reminiscent of the 2 off states recently observed spectroscopically and structurally in rsEGFP2 at RT ^35,36^.

**Figure 6.**
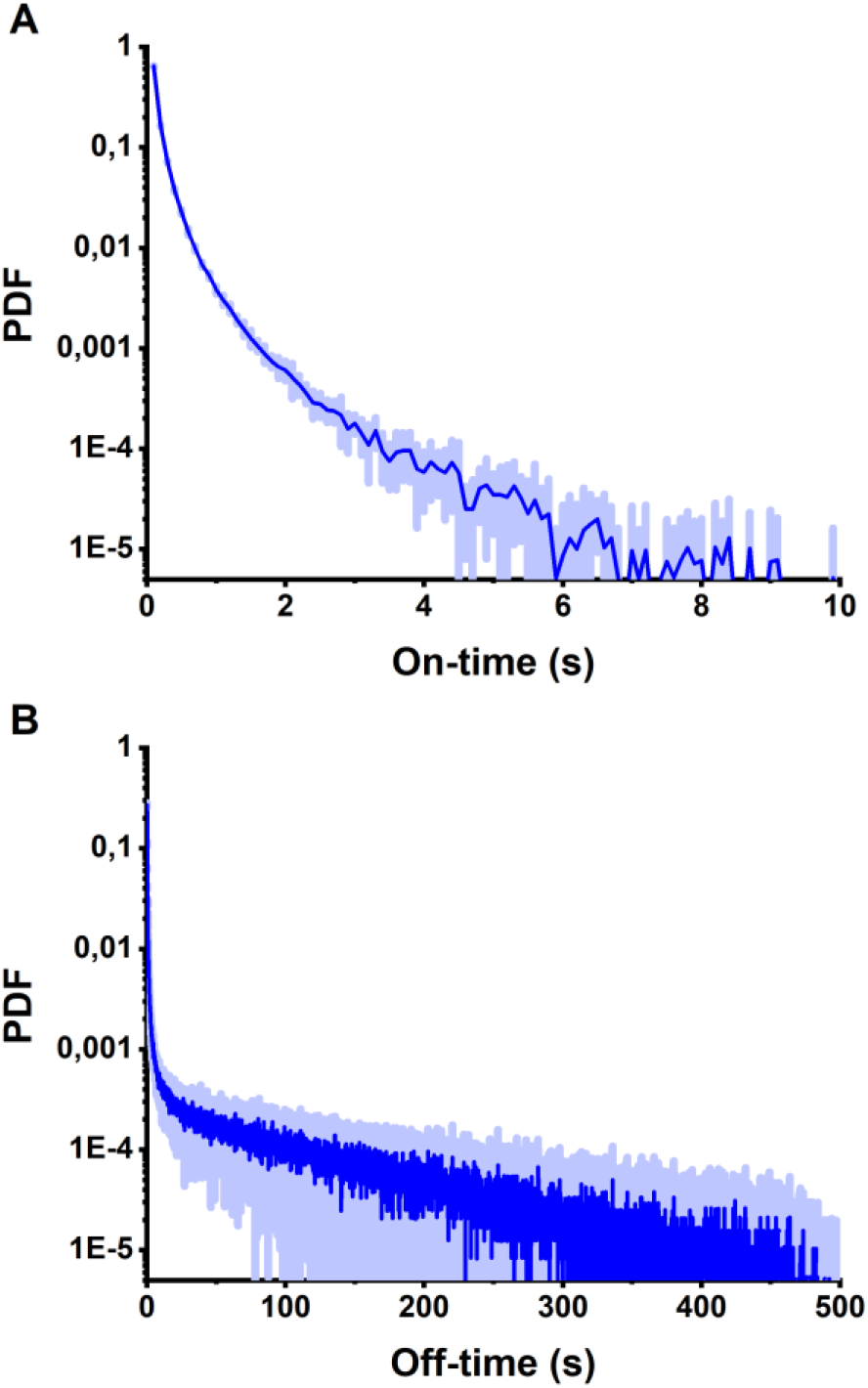
Single molecule behavior of rsEGFP2 at CT. (A) ontimes and (B) off-times histograms. The mean ± s.d. of n = 7 measurements is shown. PDF: probability density function.

The 355 nm laser we used was a nanosecond YAG laser pulsed at 3.3 kHz, so that we wondered whether pulsing played a role in the improved recovery level. The temporal profile of laser illumination would be expected to act on the switching kinetics in case of a mechanism involving 2 photons or more. Yet, titration of the on-switching rate as a function of the employed 355 nm laser power density suggested, as for 488 nm and 405 nm lasers, a single-photon mechanism (Supplementary Fig. 7A). In addition, varying the intensity of the 355 nm illumination also did not have a significant effect on the recovery level (Supplementary Fig. 7B). These data indicate that the enhanced recovery of the rsEGFP2 on-state relative to that observed by 405 nm illumination is mostly due to the blue shifted wavelength and not to the pulsed pattern of the employed 355 nm laser.

**Figure 7.**
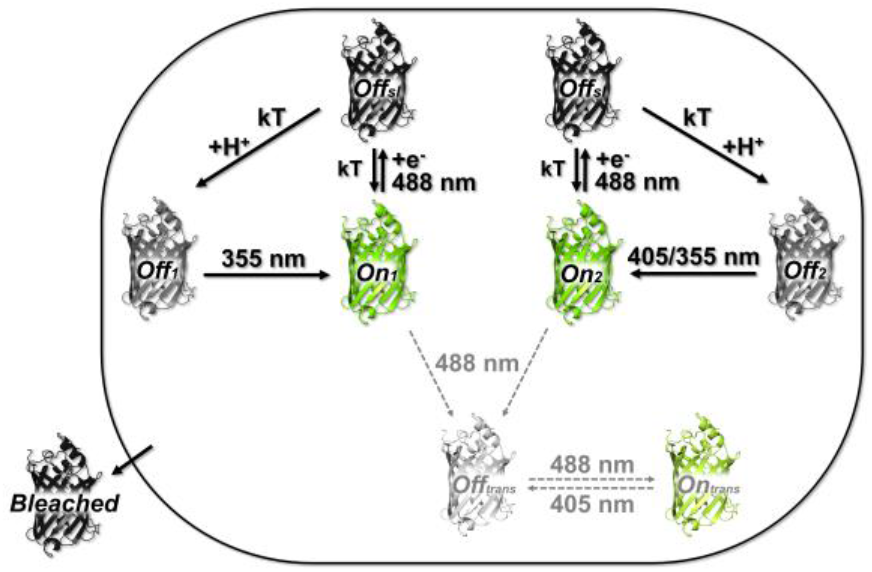
Proposed model of rsEGFP2 photophysics at CT. Sensitivity to light at defined wavelengths is indicated. kT: thermal activation. Dashed lines indicate residual involvement of *trans* chromophore states in CT-photoswitching.

Strikingly, we also observed that to achieve on-switching at similar rates, the needed 355 nm average power density was ∼20 times lower than that required using 405 nm light. Photoswitching efficiency depends on the product of the extinction coefficient by the switching quantum yield at the used wavelength. Attempts to obtain pure absorption spectra of *Off*_*1*_ and *Off*_*2*_ by computing difference spectra (Supplementary Fig. 8) suggested that, for both off states, differences in extinction coefficients at the 2 illumination wavelengths are unlikely to explain the drastic changes in switching efficiency. This suggests that, at CT, the on-switching quantum yield at the lower-energy 405 nm wavelength is significantly lower than at 355 nm for *Off*_*2*_, and practically zero for *Off*_*1*_.

**Figure 8.**
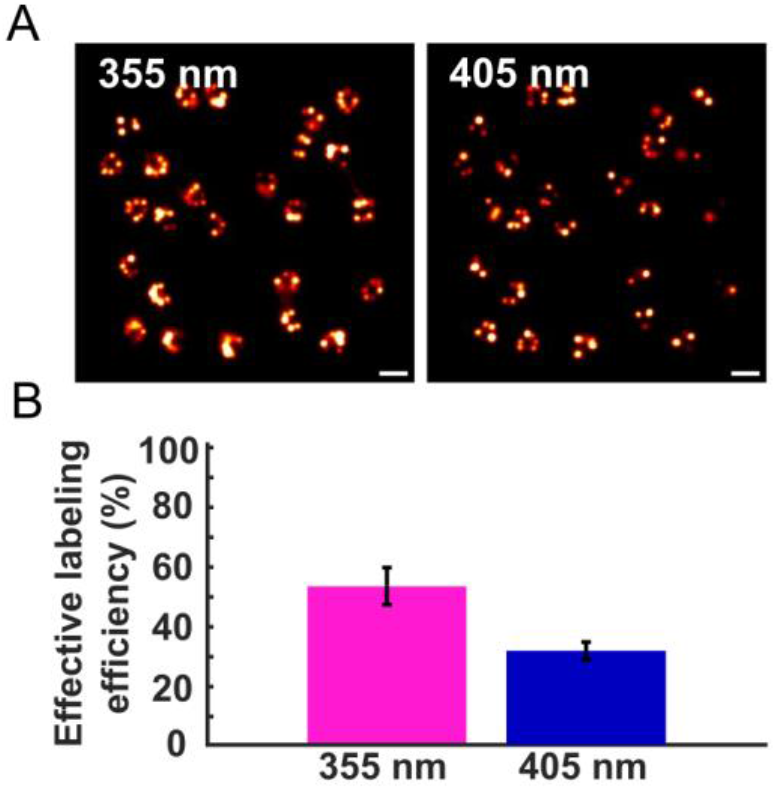
Simulated cryo-SMLM experiments of quantitative imaging of NPCs. (A) Reconstructed images using 355 nm or 405 nm illumination. (B) Effective labeling efficiency using 355 nm or 405 nm illumination. The mean ± s.d. of n = 4 simulations is shown. Scale bar: 200 nm

Off-switching of RSFPs at RT typically involves protonation of the chromophore at the hydroxybenzylidene ring. To evaluate the possible involvement of proton transfer during off-switching at CT, we measured the pH-dependence of the switching rate in flash-cooled solution samples (Supplementary Fig. 9). The results reveal a faster switching rate at low pH, consistent with a Henderson-Hasselbalch response yielding an apparent pKa of 4.9. In view of the effective pH increase upon flash cooling, such value appears consistent with the known pKa=5.8 of the rsEGFP2 chromophore at RT.

Finally, we recorded multi-switching absorbance and fluorescence curves with on-switching either induced by 405 nm or 355 nm light (Fig. 4, Supplementary Fig. 10). Photofatigue at CT was much higher than that measured at RT ^32^. This observation is in line with the incomplete recovery observed in Fig. 2A and 2B attributed in part to photobleaching by 488 nm, 405 nm and 355 nm light. Of note, photofatigue developed faster when 355 nm light was employed. Thus, the advantage of a higher on-state recovery using 355-nm light is progressively offset by faster photobleaching upon repeated switching. This finding suggests that the *Off*_*1*_ dark state is more prone to photobleaching than *Off*_*2*_, but may also relate to enhanced photodamage by UV light as compared to 405 nm light on biological material, including at CT ^41^.

In order to investigate the structural signature of the rsEGFP2 cryo off-switched states, we illuminated rsEGFP2 crystals maintained in the ∼110 K nitrogen gas stream of our microspectrophotometer with 488 nm light.

We verified by absorption microspectrophotometry that the same off-states were produced as in solution samples (Supplementary Fig. 11), and collected cryo-crystallographic data after transfer to an ESRF synchrotron beamline. No significant *cis-trans* isomerization could be observed (Fig. 5), as opposed to control measurements where crystals were illuminated at RT (Supplementary Fig. 12).

Furthermore, at the resolution of our data (2.4 Å), difference electron density maps did not allow to identify any significant conformational change between the off-switched and on-switched states, nor, consequently, between *Off*_*1*_ and *Off*_*2*_. Residual negative electron density at the level of the conserved Glu223 (rsEGFP_2_ amino-acid numbering) suggested partial decarboxylation of this residue. Decarboxylation through electron transfer via a Kolbe mechanism has previously been observed in fluorescent proteins to induce photoactivation ^31^ or photobleaching ^42^. We tentatively assign the decarboxylation observed here to photobleaching. Overall, the crystallographic data are consistent with the notion that the rsEGFP_2_ cryo off-states are predominantly *cis* states.

We also observed that chromophores switched off at RT by 488 nm laser light and then irradiated at CT with 405 nm laser light essentially stayed in the *trans*-conformation (Supplementary Fig. 13. This is in line with the observation by spectroscopy that on-switching at CT of the RT-off-switched chromophore is limited to weak recovery of a redshifted on-state (Fig. 1D). The data are in fact consistent with a scenario in which this residual red shifted on-state would originate from anionic chromophores in the *trans* configuration. Of note, significant negative difference electron density is visible on Glu223 as well as on the hydroxybenzylidene moiety of the chromophore. Decarboxylation of Glu223 in the *trans* isomer form of the chromophore might induce chromophore destabilization, a signature for photobleaching that might have resulted from the extensive 405 nm illumination employed in this experiment.

To complement our results at the ensemble level, we collected single-molecule fluorescence traces from purified rsEGFP2 molecules flash frozen on coverslips, using a PALM microscope operating at CT (110 K) (Supplementary Methods) ^21^. The flash-frozen samples were initially off-switched by 488 nm light, and 405 nm light was then turned on to elicit single-molecule activation. A representative single-molecule pattern and a single-molecule trace are shown in Supplementary Fig. 14. Analysis of the data allowed generating histograms revealing the rsEGFP2 single-molecule photophysical behaviour at CT (Fig. 6, Supplementary Fig. 14).

The on-time and off-time histograms are clearly multiphasic. The fast phases in both histograms suggest that the rsEGFP_2_ molecules rapidly toggle between an on-state (half on time = 36.5 ms) and a short-lived dark state (half off-time = 63 ms), that we refer as *Off*_*sl*_. Due to its short duration, *Off*_*sl*_ is not apparent in the ensemble data but plays a central role in the observed blinking pattern at the single-molecule level. The slow phase of the on-time histogram is mostly attributed to the fixed dipole orientation of the fluorescent proteins at CT, generating an anisotropic response to illumination (as for the kinetic traces measured at the ensemble level). The slow phase of the off-time histogram is, at least in part, attributed to the long-lived *Off*_*2*_ dark state. Molecules in *Off*_*1*_ do not contribute to the off-time histogram in the absence of 355 nm light. Of note, the median number of detected photons per merged localization (370±16 ph, Supplementary Fig. 15 is similar to typical values recorded from SMLM datasets collected at RT with green-to-red photoconvertible fluorescent proteins (PCFPs) such as mEos4b (397±42 ph, Supplementary Methods). This can be explained by the strongly increased fluorescence quantum yield of FPs at CT, which compensates for the lower numerical aperture (NA) of the air-objective used in our cryo-microscope, while in both cases, short-lived blinking interrupts photon emission by single molecules. Yet, due to the larger point spread function of the cryo-microscope, the median localization precision achieved at CT (38±0.5 nm) is lower than that at RT with PCFPs (24±3.5 nm) (Supplementary Fig. 15). Thus, with current state-of-the-art methodology, and in view of the relatively limited number of switching cycles achievable by single molecules (Fig 4), FP-based CT-SMLM data may not provide nanoscale images of superior quality as compared to RT-SMLM data.

## DISCUSSION

Combining the ensemble and single-molecule data presented above allows drawing a consistent model of the cryo-photophysical behavior of rsEGFP_2_ (Fig. 7). We propose that rsEGFP2 in its thermally relaxed on-state adopts two conformations (*On*_*1*_ and *On*_*2*_) that are in rapid exchange at RT, but that do not exchange significantly anymore at CT. These two populations may differ by e.g. different H-bonding patterns around the chromophore. *On*_*1*_, upon illumination by 488 nm light at CT, produces the non-fluorescent state *Off*_*1*_, while *On*_*2*_ switches to *Off*_*2*_. Upon illumination with 405 nm light, *Off*_*2*_ is able to switch back to *On*_*2*_, while *Off*_*1*_ remains essentially unresponsive. Upon illumination with 355 nm light, both *Off*_*1*_ and *Off*_*2*_ are able to switch back to their respective fluorescent on-states, causing more extensive recovery than with 405 nm illumination. The two populations of rsEGFP2 molecules do not photobleach at the same rate, with *On*_*1*_*/Off*_*1*_ being more prone to photodestruction. Finally, the single-molecule data revealed that *On*_*2*_, in addition to switching to *Off*_*2*_, efficiently switch to one (or several) additional shortlived dark state(s), *Off*_*sl*_, upon illumination with 488 nm light at CT. Although *On*_*1*_ was essentially unobserved in our single-molecule data (collected in the absence of 355 nm light), *Off*_*sl*_ might also be reached from this state.

We propose that, in addition, a residual fraction of the onstate rsEGFP_2_ molecules is still able to photo-isomerize to a *trans* configuration, producing a protonated off-state similar to that populated at RT. This RT-like off-state is able to deprotonate upon absorption of a 405 nm photon, producing a fluorescent *trans* state with red-shifted absorption and fluorescence-emission. This mechanism likely explains why, in cryo-PALM imaging, prior illumination of the sample at RT followed by flash cooling and single-molecule data collection at CT still elicits single-molecule blinking ^13^. However, in view of the low signal recovered in such case (Fig. 1E), this procedure likely results in very unfavorable effective labeling efficiency and should not be recommended. The RT-like off-state, with its absorption band peaking at around 400 nm, is only weakly sensitive to 355 nm light, in contrast to 405 nm light. Thus less red-shifted fluorescent molecules are produced upon 355 nm light illumination.

Radical species are known to form in FPs and are typically short-lived ^43^. We propose that *Off*_*sl*_ be an anionic or cationic short-lived radical formed via photoinduced electron transfer. We deem unlikely that *Off*_*sl*_ be the triplet state, in view of its duration (∼60 ms) that largely exceeds the triplet state lifetime (∼1 ms) measured in fluorescent proteins, albeit at RT ^44^.

The exact natures of *Off*_*1*_ and *Off*_*2*_ remain to be determined. Whether they are also radicals and reached through the triplet state would be important to know, to elucidate the main mechanism of cryo-off switching. The formation of long-lived radical species could be investigated by electron paramagnetic resonance (EPR), if sufficient sample quantities could be produced. The dependence of *Off*_*1*_ and/or *Off*_*2*_ formation on addition of triplet state quenchers, oxidizing or reducing agents would unfortunately be difficult to study due to the lack of diffusion of such molecules at CT. However, attaching quenchers such as azobenzene directly to rsEGFP2 could be an exciting alternative strategy ^45^. The pH-dependence of the rsEGFP_2_ cryo off-switching rate, on the other hand, indicates that conversion to *Off*_*1*_ and/or *Off*_*2*_ is facilitated at acidic pH lower than the chromophore pKa. This suggests a possible role of H-bonding patterns surrounding the chromophore and proton transfer during off-switching at CT, despite reduced H^+^ diffusion below the glass transition temperature ^46^. Overall, an appealing scenario is that *Off*_*1*_ and/or *Off*_*2*_ would form upon rare protonation of *Off*_*sl*_ and thus be protonated radical species. Such species might be similar to the radical state previously identified in the bi-photochromic fluorescent protein IrisFP ^47,48^.

The existence of two rsEGFP_2_ on states that would exchange at RT but not at CT, although consistent with all our data, is not fully demonstrated by our study. Yet, it is interesting to relate this hypothesis with the fact that multiple switching pathways have also been identified at RT ^36,37^. In the future, it could be interesting to investigate the cryo-switching mechanisms of the rsEGFP2 V151A and V151L mutants, which have been shown to abrogate off-state heterogeneity at RT ^36^.

Another finding of the present study is the much higher back switching quantum yield of *Off*_*1*_ by 355 nm light (× ∼20) as compared to 405 nm light. Whereas species-specific quantum yields are generally assumed to be wavelength independent, a wavelength-dependence of some reaction yields has been measured in organic molecules ^49^. However, to our knowledge, such a strong wavelength-dependence of a phototransformation quantum yield in a fluorescent protein has not been reported to date. This dependence is likely a function of temperature, exacerbated at CT. We propose that at CT, vibrational relaxation may be slowed down to a point where phototransformation pathways efficiently compete with relaxation. Thus, if photoswitching is promoted from a vibrationally excited state preferentially reached by 355 nm excitation, the corresponding quantum yield may effectively be greatly enhanced.

To evaluate the potential gain in using 355 nm rather than 405 nm light in cryo-SMLM experiments, we performed simulations using the recently developed SMIS software ^50^. Our main goal was to evaluate whether the more efficient on switching provided by 355 nm illumination could improve the effective labeling efficiency of rsEGFP_2_ despite more pronounced photobleaching. First the ensemble switching behaviour was reproduced by implementing the rsEGFP2 photophysical model described above (neglecting the residual formation of *trans* chromophores). The spectroscopic signatures of all photoactive states were included and phototransformation quantum yields (Supplementary Table 1) were adjusted to match the observed photoswitching rates as well as the differential photofatigue behavior measured with 405 nm and 355 nm light (Supplementary Fig. 16. The refined photophysical model was then used in virtual cryo-SMLM experiments aimed at quantitative imaging of the Nup96 nucleoporin within nuclear pore complexes (NPCs) ^51^. The results, presented in Fig. 8, show that the effective labeling efficiency can be raised from ∼32% to ∼54%. Thus, based on our rsEGFP_2_ photophysical model, we conclude that the more efficient recovery level offered by the 355 nm laser prevails over the more stringent photobleaching induced by this laser. This finding will now need to be confirmed in genuine experimental conditions.

## CONCLUSION

This work presents the first in-depth investigation of the cryo-switching mechanism of a fluorescent protein. We have shown that rsEGFP2, an efficient RSFP that switches at ambient temperature through coupled *cis-trans* isomerization and protonation of its chromophore, adopts a different switching mechanism at cryogenic temperature. Because of the lack of conformational freedom below the glass transition temperature (∼ 180 K), isomerization of the rsEGFP2 chromophore is largely hindered at ∼110 K. In contrast, this FP populates two off-switched states adopting *cis* configurations and displaying absorption bands largely blue shifted relative to that of the *trans* protonated off-state reached at RT. A 405 nm laser classically used in RT-SMLM switches back only one of the off-states. In contrast, a 355 nm laser applied at very low power is able to efficiently reactivate both off states and thus a larger fraction of the molecules. Cryo-SMLM simulations on the Nuclear Pore Complex suggest that indeed there would be a net gain in using such a laser for real studies thanks to an increase of effective labeling efficiency. Yet, cellular damage induced by 355 nm light, although likely limited at cryogenic temperature, would need to be evaluated. It will also be interesting in the future to evaluate whether the results reported here can be generalized to other FPs such as mEmerald ^14^. As the cryo switching mechanism of rsEGFP2 does not seem to relate to the switching mechanism adopted at RT, we anticipate that FPs that are not necessarily phototransformable at RT may share the described behaviour and be sensitive to reactivation by 355 nm light. In fact, the various FPs studied by Tuijtel et al ^13^ appear to show relatively similar behaviors, including EGFP. Of note, in the case of Padron, it is remarkable that both negative and positive cryo switching have been described ^13,27^. It is likely that this protein, while it can sustain *trans* to *cis* isomerization at CT (positive switching) can also undergo the same cryo-switching mechanism as rsEGFP2 (negative switching). The present study highlights that population heterogeneity is a major hallmark of fluorescent protein photophysical behavior. Removing this heterogeneity in favor of only the *On*_*2*_ population would be an even more attractive option than using 355 nm light for efficient activation of rsEGFP2 at CT. In conclusion, the present study opens the door to FP engineering and optimization of illumination conditions to improve cryo-photoswitching in cryo-SMLM.

## ASSOCIATED CONTENT

**Supporting Information**. Supplementary Methods, Supplementary Figures and Supplementary Tables. This material is available free of charge via the Internet at http://pubs.acs.org.

## AUTHOR INFORMATION

### Author Contributions

D.B. conceived and supervised the project. A.M., O.G. and D.T. performed microspectrophotometry experiments. O.G. performed single-molecule experiments. A.M. and V.A. performed crystallography experiments. J.W. performed simulations. A.M., V.A., J.W. and O.G. processed the data. D.T. discovered the advantage of using a 355 nm laser. M.B. and O.G. developed and optimized the microspectrophotomer instrument. I.G., O.N. and J.E. designed the cryogenic super-resolution microscope. D.B., O.G., V.A. J.W. and A.M. wrote the manuscript with input from all authors.

## ACKNOWLEDGMENT

We thank Antoine Royant for the loan of the 355 nm laser and Ninon Zala for rsEGFP2 protein production. Michel Sliwa is acknowledged for providing insight on wavelength-dependent phototransformation quantum yields.

## ABBREVIATIONS

SMLM: single molecule localization microscopy;
(d)STORM: stochastic optical reconstruction microscopy;
PAINT: point accumulation in nanoscale topography;
PALM: photoactivation localization microscopy;
RT: room temperature;
CT: cryogenic temperature;
FPs: fluorescent proteins;
RSFPs: reversibly switchable fluorescent proteins;
PCFPs: photoconvertible fluorescent proteins;
CLEM: correlative light and electron microscopy;
EPR: electron paramagnetic resonance;
ESRF: European Synchrotron Radiation Facility;
NPC: nuclear pore complex.

