## Supporting Information for "Photophysical studies at cryogenic temperature reveal a novel photoswitching mechanism of rsEGFP2"

### Supplementary Methods

#### *Protein expression and purification*

rsEGFP2 was expressed and purified as described earlier.[1] Briefly, expressing cells were lysed and 6xHis-tagged rsEGFP2 was purified by gravity flow immobilized metal affinity chromatography (IMAC) using 100 ml of Ni-NTA resin, followed by size exclusion chromatography on a HiLoad 16/600 Superdex 75 prep grade column.

#### *Solution sample preparation*

For microspectrophotometry experiments at pH 7.5, purified rsEGFP2 was mixed with a buffer solution containing 50 mM HEPES pH 7.5, 20% glycerol and 50 mM NaCl, providing a final protein concentration of 20 mg/ml. For pH-dependent experiments, purified rsEGFP2 was mixed with buffer solutions containing 20% glycerol and 1M MES (pH5 and 6), 1M HEPES (pH7) or 1M Tris (pH9), providing a final protein concentration of 7 mg/ml.

#### *Solution sample illumination*

For microspectrophotometry experiments at RT, rsEGFP2 solution samples were sucked into capillaries of 0.5 mm x 1 mm section attached to a goniometer base. The capillaries were then mounted to our home-built microspectrophotometer.[2]

For microspectrophotometry experiments at CT, solution samples were prepared in the same way, but flash-cooled in a 110 K nitrogen gaseous cryo-stream (Oxford Cryosystems) connected to the microspectrophotometer.

Samples were illuminated with alternating 488 nm and 405 nm or 355 nm lasers for on-to-off and off-to-on protein switching, respectively, according to the scheme shown in Supplementary Fig. 17. During on-to-off switching phases, absorption and fluorescence spectra were recorded in an interleaved manner. During off-to-on switching phases, only absorption spectra were collected to avoid unwanted off-switching by 488 nm excitation light.

The cycle time was 1 second. Laser illumination, white-light illumination and spectroscopic data collection were synchronized using a digital delay-pulse generator (9518, Quantum Composer). For on-to-off switching, the pulse sequence started with triggering of a Deuterium-Halogen lamp (AvaLight-DH-S-BAL, Avantes) and one of two CCD-based spectrometers (AvaSpec ULS 2048L-USB2, Avantes) to record absorption spectra (integration time of 30 ms with averaging of 5 spectra). Then a fiber coupled 488 nm laser (LBX-488-200, Oxxius) was used to induce rsEGFP2 off-switching for 750 ms (75% of the cycle time). Fluorescence spectra were collected by recording fluorescence emission excited by the 488 nm laser in an epifluorescence mode for 1 ms, 5 ms after the start of laser illumination. Emitted light passed through a 488 nm dichroic mirror (Di02-R488, Semrock) and a notch filter (NF03-488E-25, Semrock), and was detected by the second AvaSpec spectrometer, using an integration time of 1 ms. For off-to-on switching, absorption spectra were collected using the same protocol except that a fiber coupled 405 nm laser (LBX-405-200, Oxxius) or a 355 nm laser (MPL-F-355, CNI) were used with a reduced duty cycle (30% of the cycle time), starting after 600 ms. Pulsing frequency of the YAG-based 355 nm laser was measured to be 3.3 kHz, with pulse durations of ~10 ns.

Laser powers were measured at sample position using a calibrated power meter (PM100D, Thorlabs) with a 355 nm light sensitive sensor (S170C, Thorlabs) and another sensor sensitive to 488 nm and 405 nm light (S121C). Laser beam profiles (circular top flat) were recorded from images collected through the objective of the microspectrophotometer onto a CCD camera. Laser power densities were calculated by dividing the measured powers by the beam profile surface areas.

At CT, using input optical fibers of 200 µm diameter (25 µm spot size at sample position) laser power densities were adjusted in the range 430-5700 W/cm² (488 nm), 120-780 W/cm² (405 nm ) and 4-33 W/cm² (355 nm). At RT, laser power densities of 81 W/cm² (488 nm) and 4 W/cm² (405 nm) were used, except for Supplementary Fig. 1, where similar laser power densities as used at CT were employed (0.5 kW/cm² and 0.1 kW/cm² for 488 nm and 405 nm lasers, respectively).

#### *Crystal preparation*

rsEGFP2 crystals were grown at 20°C by the hanging drop vapor diffusion method with a drop of 2 µl of rsEGFP2 solution at 13 mg/ml mixed with 2 µl mother liquor against a 1 ml mother liquor solution containing 100 mM HEPES buffer, pH 8.1 mixed with 1.9 M ammonium sulfate. Crystal dimensions were ~50 × 40 × 200 µm³. Crystals were harvested with cryo-loops, soaked for a few seconds in a cryo-protectant solution containing the mother liquor mixed with 20% glycerol, and immediately flash cooled.

#### *Crystal pre-illumination*

Crystals were illuminated under continuous cooling (110K) on our home-built micro-spectrometer, as described in the *Solution sample illumination* section.

Crystal 1 (off-switching at CT, Supplementary Table 2) was left non-illuminated on ~half of its volume to allow collecting the structure of the on state as a control (Position 1). The other half was illuminated with the 488 nm laser (600 µm fiber diameter, ~80 µm spot size) for 700 s to allow collecting the structure of the cryo-switched off state (Table 2, Position 2). The laser power density was kept at low level (46 W/cm²), to preserve the crystal from heating and partial disordering, considering the very high optical density of rsEGFP2 crystals in the on-state at 488 nm. Progression of off-switching was monitored by fluorescence and absorption microspectrophotometry (Supplementary Fig. 12) using the same acquisition scheme described in the *Solution sample illumination* section.

Crystal 2 (off-switching at RT, back on-switching at CT, Supplementary Table 3) was switched off at RT over its whole volume with 488 nm laser light (3.4 W/cm², 600µm fiber) for 30 s directly in its crystallization drop. The crystal was then flash cooled and mounted on the microspectrophotometer, and illuminated at 110K on ~half of its volume with the 405 nm laser (14W/cm², 600 µm fiber diameter, ~80 µm spot size) for 180 s. Progression of residual on-switching was monitored by absorption microspectrophotometry (Supplementary Fig. 13B) using the same acquisition scheme described in the *Solution sample illumination* section. The non-405 nm illuminated part of the crystal was used to obtain the structure of the RT-switched off state (Table 3, Position 1) and the illuminated part was used to obtain the RT-off-switched and then cryo-back-switched state (Table 3, Position 2). The extensive exposure to 488 nm light at RT, used to ensure complete off switching of the crystal, possibly led to partial photobleaching.

The on-state structure obtained from Crystal 1 (Position 1) and the RT-switched off-state structure obtained from Crystal 2 (Position 1) are shown together in Supplementary Fig. 12 to confirm the observation of a *cis-trans* isomerization at RT with our setup.

#### *Crystallographic data collection*

X-ray diffraction data sets were collected at 100 K at the European Synchrotron Radiation Facility (ESRF, Grenoble, France) on the beamline ID30A-3/MASSIF-3 (Crystal 1) equipped with an Eiger_4M (Dectris) detector and on the beamline ID30B of the ESRF (Crystal 2) equipped with a Pilatus3_6M (Dectris).

#### *Structure Refinement*

Data was processed with AutoProc[3] (Crystal 1) and XDSAPP[4] (Crystal 2), and merged data were phased by molecular replacement with MOLREP,[5] within the CCP4i2 suite,[6] using the rsEGFP2 on-state structure (PDB entry 5DTX) as starting model for Crystal 1 and the rsEGFP2 off-state (PDB entry 5DTY) as a search model for Crystal 2. To obtain the final models, refinement was performed with Refmac5[7] in consecutive cycles (with isotropic B-factors) and finalised with Phenix[8] and manual model building in-between cycles with Coot.[9] As expected, the on-state structure of Crystal 1 was successfully fitted with the model in cis-conformation (PDB entry 5DTX). As for the off-state of Crystal 1 refinement with the chromophore in trans-conformation was not successful and the chromophore appeared to be in the *cis*-conformation as well, we used the cis-conformation model also for the off-state which turned out to be successful. For Crystal 2 both states, the off-state and the off-to-on switched state, could be only fitted successfully with the *trans*-conformational model (PDB entry 5DTY).
The structures were validated using Molprobity[10] and Coot validation tools.

#### *Analysis of kinetic evolution of spectral data*

Absorbance spectra were individually baseline corrected with a constant offset calculated based on the average optical density in the 550-650 nm range, eventually corrected to avoid negative optical densities in the 480-540 nm of the spectra. Spectra were smoothed with a moving average filter (smoothing factor of 10) using Matlab 2021a (The MathWorks). On-state populations were monitored by integrating absorbance spectra between 470 and 500 nm, or fluorescence spectra between 495 and 630 nm. To extract decay and recovery rates, off-switching and on-switching curves were fitted with bi-exponential models (ExpGro2 and ExpDec2, respectively) in Origin 2021b (OriginLab Corporation). The rates were calculated taking into account the whole measurement period of 1 s (instead of actual laser illumination times) for each data point.

For calculation of the rsEGFP2 on-state recovery levels, the difference between the maximum and minimum value of the recovery (on-switching) phase was dived by the difference between the maximum and minimum value of the initial off-switching phase.

#### *Extraction of Off1 and Off2 absorption spectra*

The absorbance spectrum of the protein in the initial on-state (Supplementary Fig. 8B, green), the spectrum after off-switching with 488 nm laser light (black), and the spectrum of the sample after back on-switching with 405 nm laser light (magenta) were analysed to extract the spectra of each off-state. The on-state spectrum (green) was first scaled to match the shape of the on-state spectrum back-switched with 405 nm light (magenta) to provide the light green spectrum in Supplementary Fig. 8B. Then this scaled spectrum (light green) was subtracted from the back-switched spectrum (magenta), providing the $Off_2$ spectrum (Supplementary Fig. 8A, blue). Finally, the $Off_2$ spectrum was subtracted from the experimental off-state spectrum (black), which contains the signatures of both off-states together, to provide the $Off_1$ spectrum (Supplementary Fig. 8A, magenta).

#### *Estimation of residual off-switching by the 405 nm laser*

Residual off-switching by the 405 nm laser was estimated using the ensemble fluorescence on- and off-switching data. First, the off and on-switching rates were estimated using the half-life and half rise time of the fluorescent intensity under 488 nm and 405 nm illumination respectively. The off and on-switching quantum yields were then calculated by dividing the rates by the respective excitation rates (off-switching QY = 5.4e-8, on-switching QY = 2.8e-5). Finally, the switching contrast was estimated by calculating the on- and off-switching rates under 100W/cm$^2$ 405-nm illumination, which gave an off-switching rate of 4.8e-4 s$^{-1}$ and an on-switching rate of 0.43 s$^{-1}$, resulting in an off:on ratio of ~900.

#### *Ensemble and single-molecule simulations*

Simulations were performed using the recently developed SMIS software (ref). A simplified photophysical model of rsEGFP2 was used consisting of two on-state populations ($On_1$ and $On_2$) with corresponding off and dark states. The photo switching quantum yields were estimated using the rates extracted from the experimental ensemble and single-molecule data. The photoswitching fatigue experiments were used to estimate the relative populations of $On_1$ and $On_2$ (70% and 30% respectively) and the photobleaching quantum yields. It should be noted that different combinations of photobleaching quantum yields from the *On* and *Off* states provide the same ensemble behavior (Supplementary Table 1 and Supplementary Fig. 16).

Single-molecule simulations were performed in 2D using nuclear pore complexes (NPCs) virtually labeled with rsEGFP2 (8 corners, 4 molecules per corner). Off-switching was achieved by 1200 W/cm$^2$ 488 illumination and on-switching by 0.01 W/cm$^2$ 355 nm or 0.2 W/cm$^2$ 405 nm light illumination. Molecules were localized using Thunderstorm,[11] after which the effective labeling efficiency was determined using SMAP.[12] Four simulations of 1 million frames with 25 NPCs each were run.

*Single-molecule measurements at cryogenic temperature*

Single-molecule measurements were performed on a home-built setup (Supplementary Fig. 18). The microscope body and the cryostat were kindly provided to us by Joerg Enderlein et al and are thoroughly described in a dedicated publication.[13]

Samples were prepared by mixing 1 µl of purified rsEGFP2 fluorescent protein and 1 µl of 1/100 stock dilution of TetraSpek beads (T7279, Thermo Fisher Scientific) in 100 µl of 0.3% w/v PVA (363138-25G, Sigma-Aldrich) in 50 mM HEPES, 300 mM NaCl buffer at pH7.5, to obtain a final protein concentration of 100 nM. Then, round fused silica cover glasses, 12 mm in diameter (Shanghai Wechance Industrial Co., Ltd), were cleaned in a UV-ozone cleaner (UVOCS) for 20 min and spin-coated with a protein/polymer mixture (7.5 µl drop) in two steps: 30 s at 1000 rpm followed by 60s at 3000 rpm.[14] The cover glass was next plunge frozen in LN2, mounted on the sample holder and transferred to the cryostat. A vacuum of ~$10^{-3}$-$10^{-4}$ Pa was regenerated with a turbo pump (HiCube, Pfeiffer Vaccum). A temperature sensor installed at the level of the sample-mount chamber gave a value of 110 K. A flexible tube delivering dry gaseous $N_2$ was taped to the bottom of the cryostat to prevent residual condensation/ ice formation during experiments.

To excite rsEGFP2, the sample was illuminated with a 488 nm laser (LBX-488-200, Oxxius) at 275 W/cm². This power density was shown in previous studies to be sufficiently low to not induce sample devitrification.[15] As an additional control, we compared the rsEGFP2 emission spectrum on the single-molecule cryo-microscope with that measured on the microspectrophotometer (Supplementary Fig. 19). Both spectra showed similar band narrowing when compared to a spectrum collected at RT.

A UV laser at 405 nm (06-MLD 405 nm, Cobolt), set to 1 mW/cm², was used for protein activation. This very small power density was used to account for the rather high protein concentration of our samples.

Fluorescence was collected by a long working distance Olympus air objective (0.7 NA) and detected by an EMCCD camera (Evolve 512, Photometrics). The pixel size measured with a Thorlabs ruler (R1L3S2P) was 276 nm/px.

The integration of a fiber-coupled 355 nm laser to the cryo-microscope is currently in progress.

Single molecule data were acquired with Micro-Manager 2.0. The camera exposure time was set to 100 ms and EM Gain to 200. The acquisition sequence was programmed in Labview, which sends trigger pulses to camera and lasers (Supplementary Fig. 20).

Single molecules were localized using the ThunderSTORM plugin of ImageJ and data was further analyzed with in-house Matlab code.[16] Active times were calculated as the sum of on- and off-times (as most molecules were not irreversibly photobleached at the end of the acquisitions). Number of photons and localization precision were exported from ThunderSTORM after merging localizations belonging to the same molecule (maximum distance between two localizations: 20 nm; blinking gap: 1). To compare the photon budget and localization uncertainty of rsEGPF2 with a standard PCFP, single-molecule data of mEos4b from previous work[16] was used. These data were processed in the same manner as the rsEGFP2 data presented in this work to extract the median photon budget per merged localization and the median localization uncertainty.

*References*

(10) Williams, C. J.; Headd, J. J.; Moriarty, N. W.; Prisant, M. G.; Videau, L. L.; Deis, L. N.; Verma, V.; Keedy, D. A.; Hintze, B. J.; Chen, V. B.; Jain, S.; Lewis, S. M.; Arendall, W. B.; Snoeyink, J.; Adams, P. D.; Lovell, S. C.; Richardson, J. S.; Richardson, D. C. MolProbity: More and Better

Reference Data for Improved All-Atom Structure Validation. *Protein Sci* **2018**, *27* (1), 293–315. https://doi.org/10.1002/pro.3330.

**Supplementary figures**

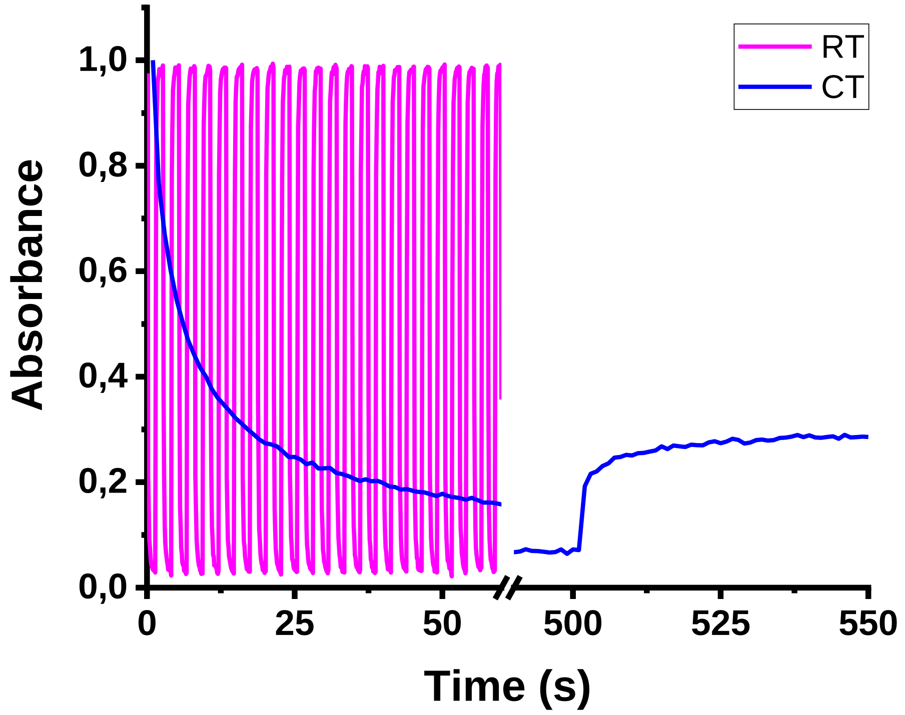


*Figure S1: Comparison of rsEGFP2 photoswitching at room and cryogenic temperature. Absorbance is measured by integration of the absorbance spectra in the 470-500 nm spectral range. For data at RT (magenta), alternative illumination at 488 nm (1.4 s, 490 W/cm²) and 405 nm (1.4 s, 100 W/cm²) was carried out 23 times. For data at CT (blue), illumination at 488 nm (5.7 kW/cm²) was performed for 500 seconds followed by illumination at 405 nm (0.95 kW/cm²) for 50 seconds. Data normalized to 1 at the start of acquisition.*

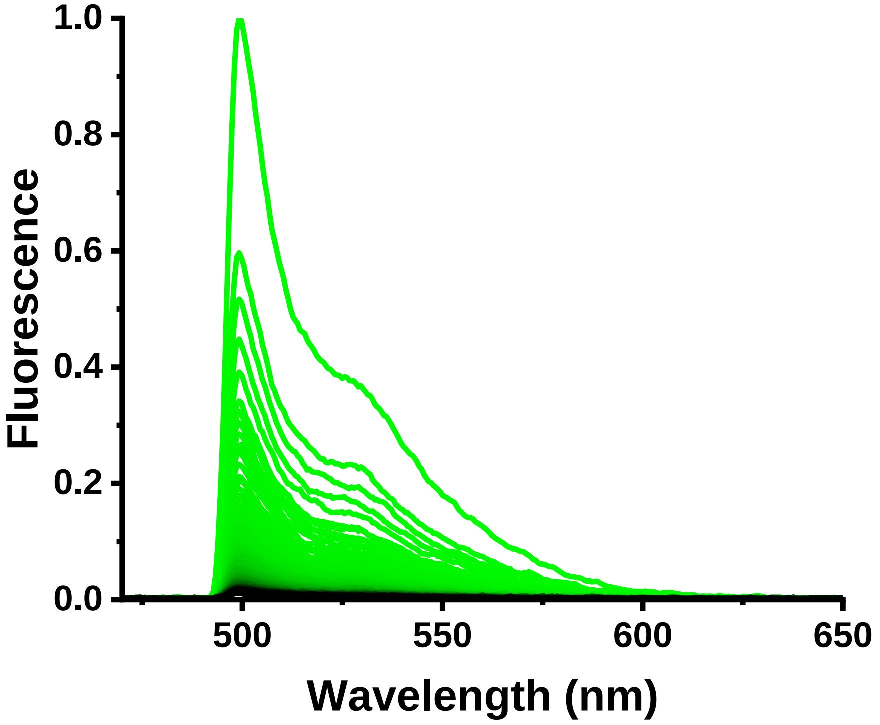


*Figure S2: On-to-off photoswitching (green-to-black) of rsEGFP2 at 110 K with 488 nm laser light (5.8 kW/cm²) applied for 500 s, monitored by fluorescence microspectrophotometry. Fluorescence emission spectra were normalized to 1 at the maximum value of the first spectrum.*

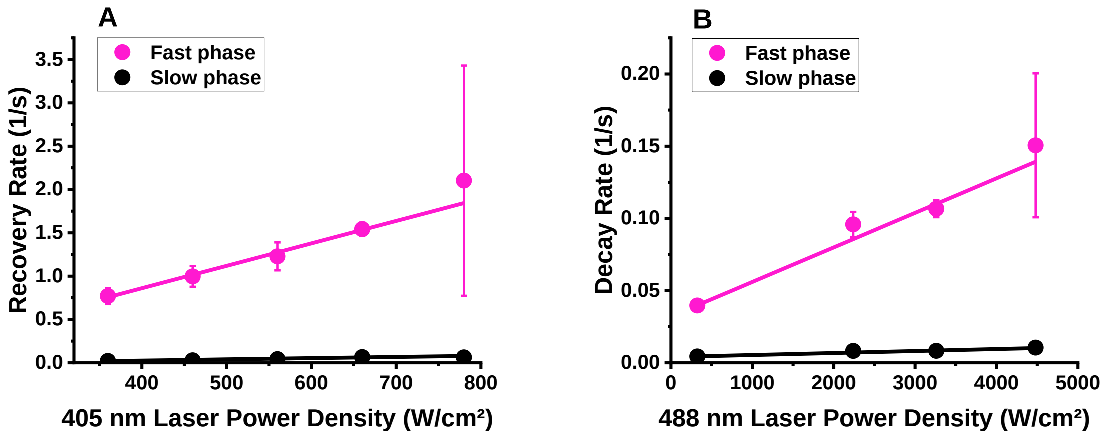


*Figure S3: Rates of rsEGFP2 off-to-on switching at CT as a function of 405 nm laser power density (A) and on-to-off switching as a function of 488 nm laser power density (B). Mean rates and standard deviations were extracted from bi-exponential fits of 3 independent switching measurements. Straight lines show linear fits to experimental rates as a function of laser power densities.*

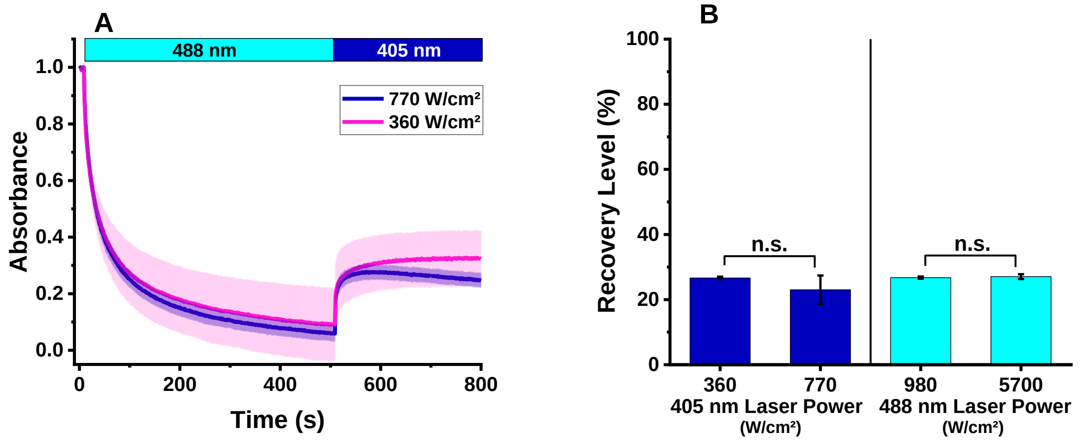


*Figure S4: (A) Evolution of rsEGFP2 recovered on-state absorbance upon illumination with medium or strong 405 nm laser light. The decay of absorbance observed at 0.8 kW/cm² suggests the onset of photobleaching. Absorbance is calculated by integration of the absorption spectra in the 470-500 nm spectral range. Absorbance switching kinetics was measured at 110 K by alternate illumination (indicated in the upper bar) with 488 nm (1.0 kW/cm²) and 405 nm laser light with lower (0.36 kW/cm²) or higher power density (0.77 kW/cm²). Absorbance is normalized to 1 at start of acquisition. The mean ± s.d. of n ≥ 3 measurements is shown. (B) Off-to-on recovery levels compared for two different 405 nm laser power densities, using 488 nm laser light at 1.0 kW/cm² for initial off switching, or two different 488 nm laser power densities, using 405 nm laser light at 0.5 kW/cm² for recovery. n.s: not significant.*

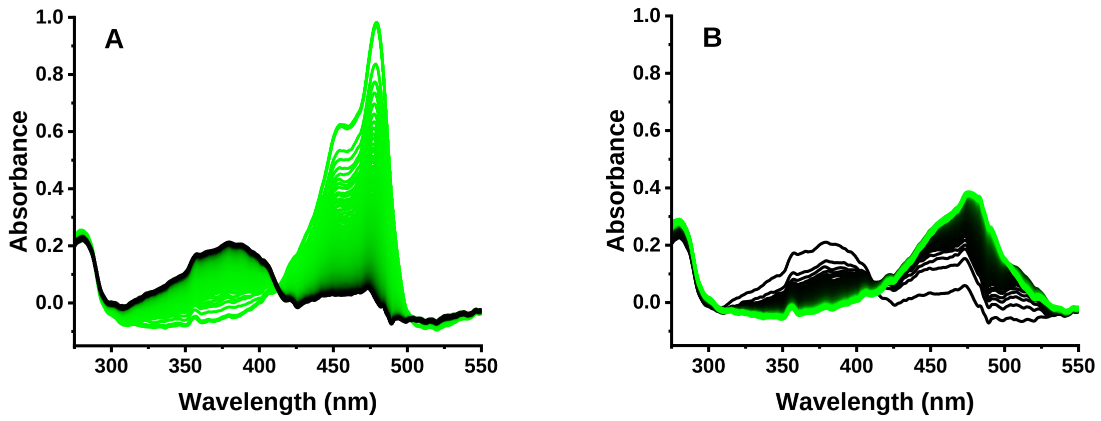


*Figure S5: On-to-off photoswitching (green-to-black) of rsEGFP2 at 110 K with 488 nm laser light (0.9 kW/cm²) for 500 s (A) and off-to-on photoswitching (black-to-green) with 355 nm laser light (0.015 kW/cm²) for 500 s (B) monitored by absorption microspectrophotometry. Spectra were normalized at the anionic chromophore peak of the first spectrum in A.*

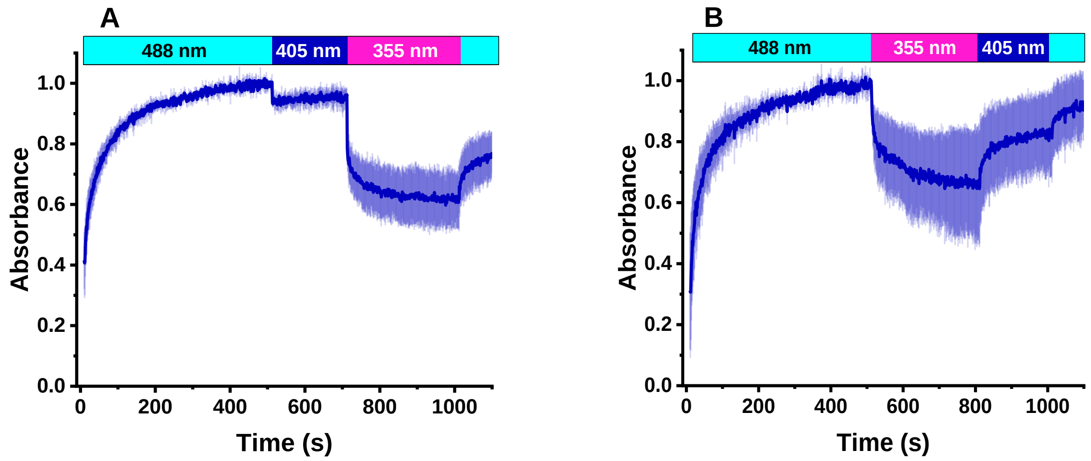


*Figure S6: Evolution of absorbance at 320 nm, representative of the Off$_1$ population, during rsEGFP2 switching at CT with 488 nm (0.4 kW/cm²), and subsequent illumination with 405 nm (0.3 kW/cm²) followed by 355 nm laser illumination (0.03 kW/cm²) (A), or 355 nm (0.02 kW/cm²) followed by 405 nm laser illumination (0.3 kW/cm²) (B), according to the schemes indicated in the upper bars. Integrated absorbance between 315 and 325 nm was measured. Absorbance was normalized to 1 at the highest value of the first phase (488 nm illumination). The mean ± s.d. of n = 3 measurements is shown.*

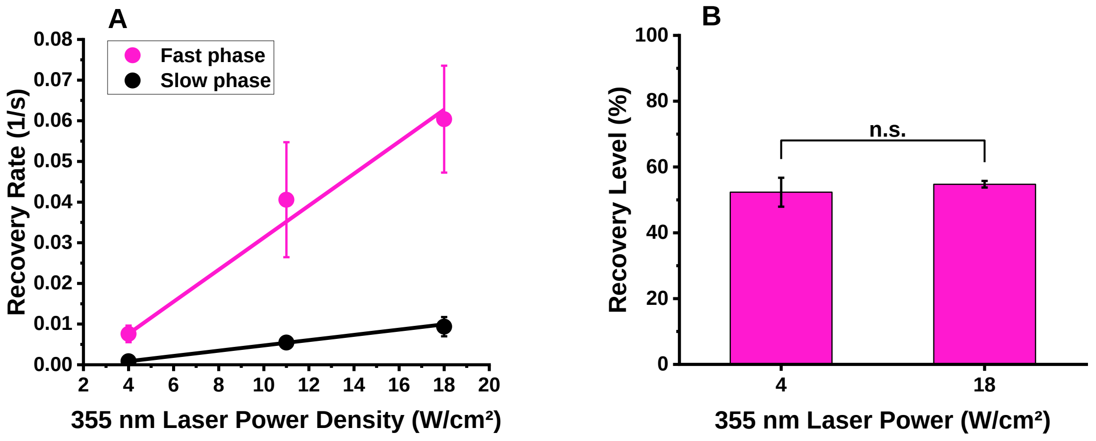


*Figure S7: (A) Rates of off-to-on switching of rsEGFP2 at CT as a function of 355 nm power density extracted from bi-exponential fits of 3 independent switching measurements. The straight lines show linear fits to extracted rates as a function of laser power density. (B) On state recovery level for two different 355 nm laser power densities. For pre-on-to-off switching, 488 nm laser light of 1.0 kW/cm² was used for all measurements. n.s: not significant.*

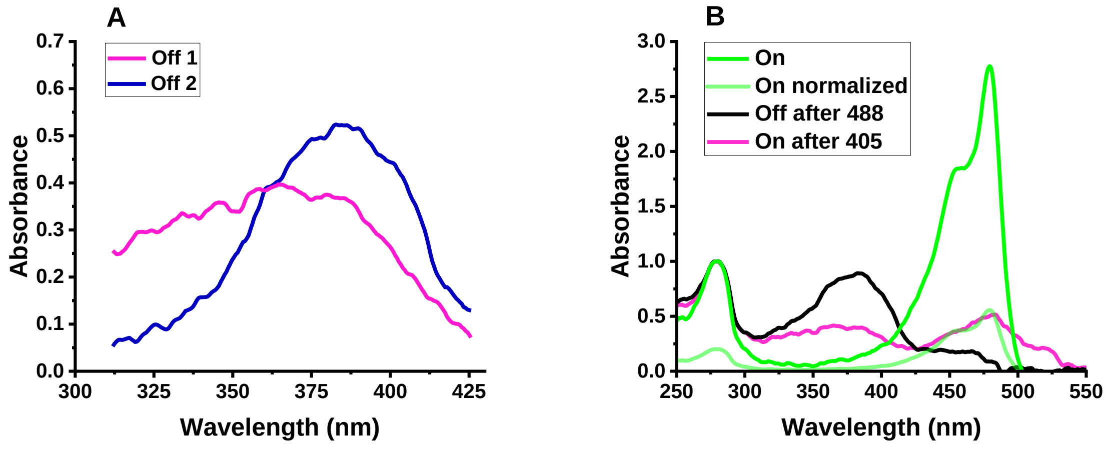


*Figure S8: (A) Estimated Off$_1$ and Off$_2$ absorption spectra extracted by computation of difference spectra. Scaling of the spectra incorporates relative populations in Off$_1$ and Off$_2$ (B) Absorption spectrum of rsEGFP2 in the on-state (green, and normalized version in light green), in the off-state (black) and in the recovered on-state after preliminary off- and on-switching (magenta). The absorption spectrum of Off$_1$ was obtained by subtracting the normalized on-state spectrum from the recovered on-state spectrum. The Off$_2$ absorption spectrum was obtained by subtracting Off$_1$ from the off-state spectrum.*

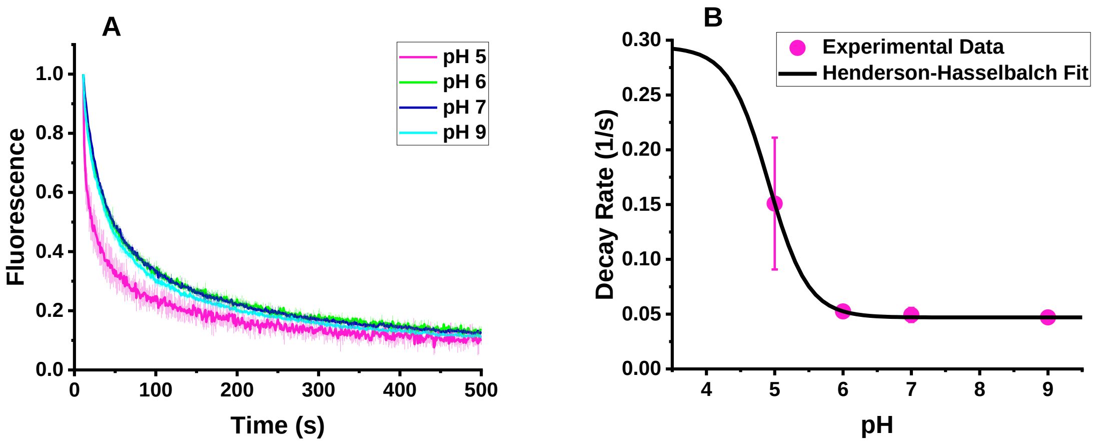


*Figure S9: pH dependence of rsEGFP2 off-switching at CT. (A) off-switching curves at different pH values (488 nm laser light, 1.0 kW/cm²). Fluorescence levels are calculated by integration of the fluorescence emission spectra in the 495-630 nm spectral range. (B) Decay rates corresponding to the fast phase of on-to-off switching, as a function of pH and extracted from bi-exponential fits of 3 independent switching measurements. The dark curve shows a fit with a Henderson-Hasselbalch function.*

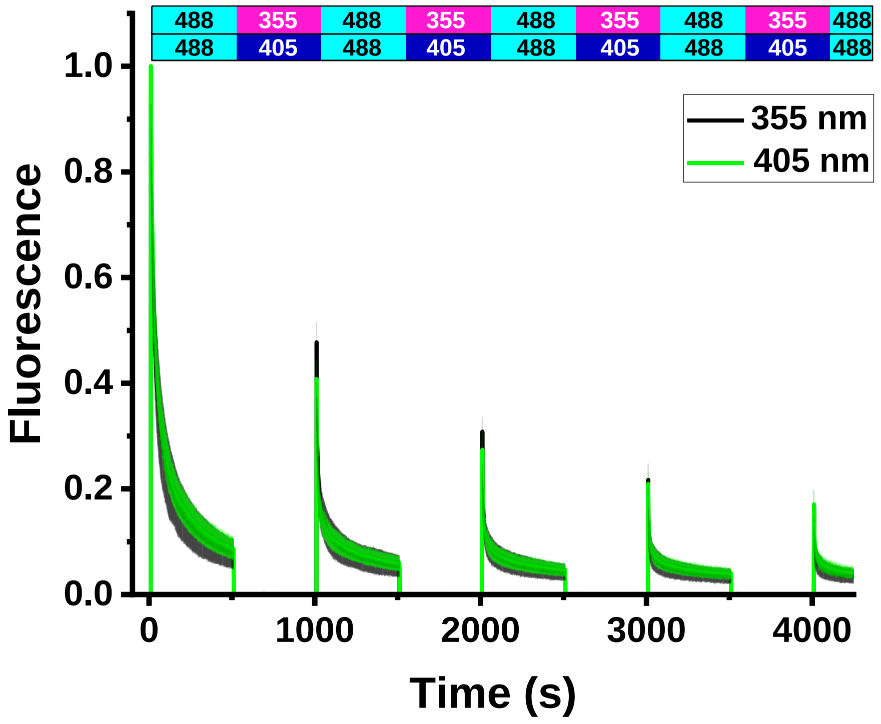


*Figure S10: Fluorescence photofatigue switching curves of rsEGFP2 at CT, corresponding to the absorbance photofatigue data shown in Fig. 4. Fluorescence was calculated by integration of the fluorescence emission spectra in the 495-630 nm spectral range. Back and forth switching was performed with 488 nm (1.0 kW/cm², green) and either 405 nm (0.2 kW/cm², green) or 355 nm (0.01 kW/cm², black) laser light, corresponding to the illumination schemes shown in the upper bars. Fluorescence was normalized to 1 at the start of acquisition. The mean ± s.d. of n = 3 measurements is shown.*

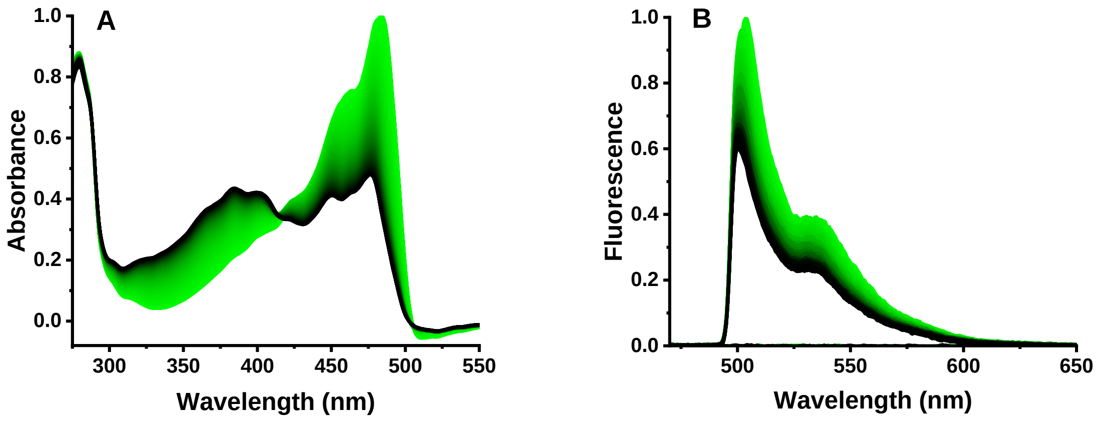


*Figure S11: On-to-off photoswitching (green-to-black) of an rsEGFP2 crystal at 110 K with 488 nm laser light (0.05 kW/cm²) for 700 s monitored by (A) absorption and (B) fluorescence emission microspectrophotometry. Spectra were normalized to 1 at the highest value of the first spectrum.*

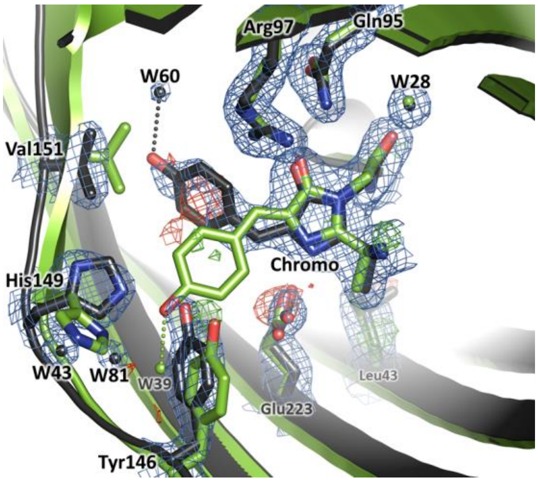


*Figure S12: Crystallographic view of rsEGFP2 switching at RT (control experiment). Refined models of the chromophore and surrounding residues of rsEGFP2 are shown in the cis on state (green carbons and water molecules) and in the trans off state (dark grey carbons and water molecules).*

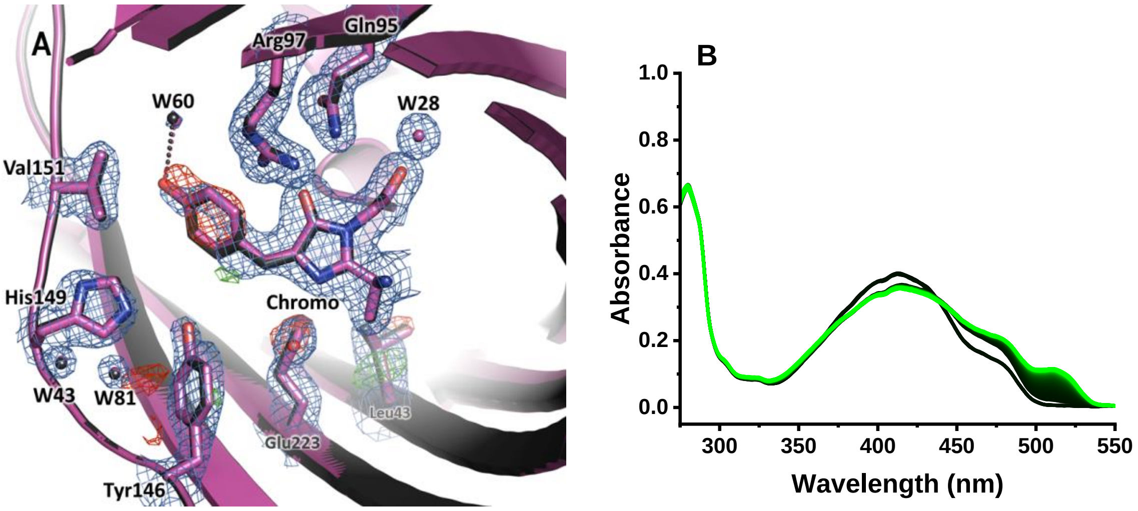


*Figure S13: (A) Refined models of the chromophore and surrounding residues of rsEGFP2 in its trans off state switched at RT by 488 nm illumination (3.4 W/cm²) (dark grey carbons and water molecules) and in the same state followed by 405 nm illumination (14 W/cm² for 180 s) at CT (purple carbons and water molecules, PDB model 8AHB). Both structures nearly perfectly overlap, so that only the second one is visible. The 2Fobs-Fcalc electron density map contoured at 1σ (blue) and the Fobs-Fcalc difference electron density maps contoured at ±3 σ (red: negative, green: positive) of the RT-switched state followed by 405 nm illumination at 110K are shown. (B) Off-to-on switching (black-to-green) by 405 nm illumination was monitored in crystalline rsEGFP2 by absorption microspectrophotometry. The absorbance spectra were normalized at the protonated peak as in Fig. 1E.*

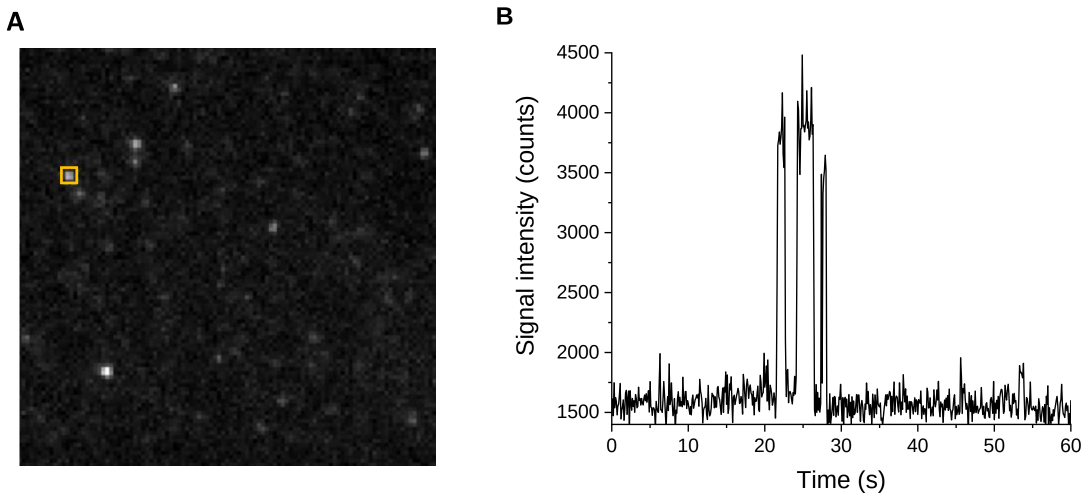


*Figure S14: Single-molecule data collection on rsEGFP2 at CT. A: example of a single frame with individual localizations of rsEGFP2 molecules. B: intensity trace during acquisition corresponding to the single molecule highlighted in yellow in A.*

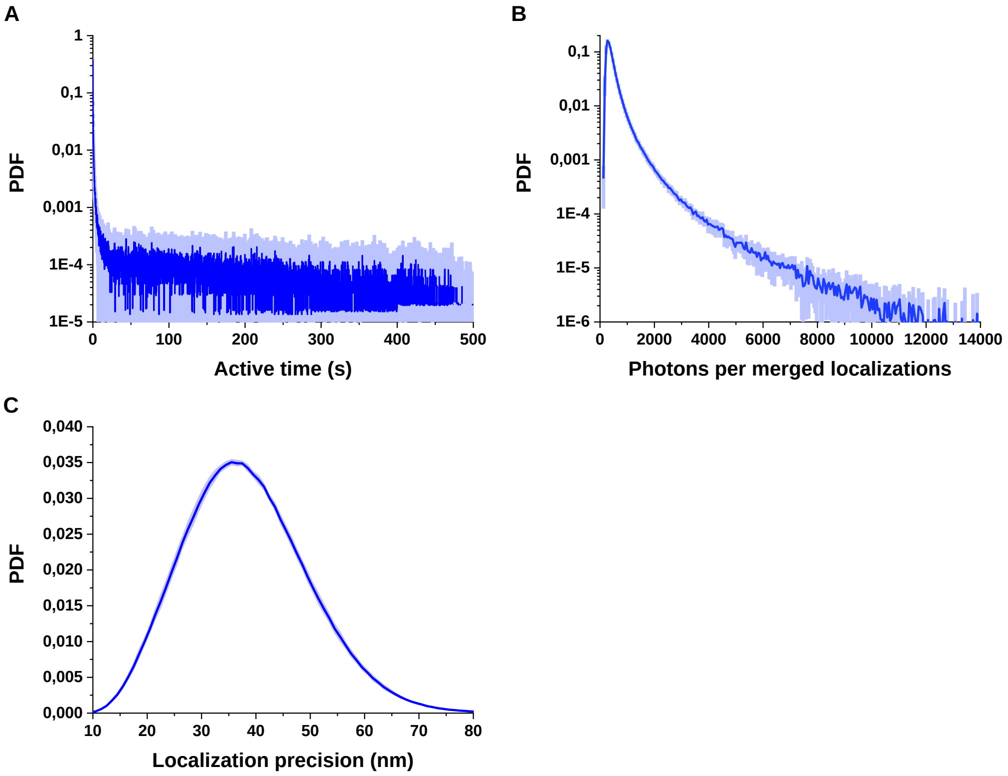


*Figure S15: (A) active time (sum of on- and off-times), (B) photon budget and (C) localization precision probability density functions extracted from CT single-molecule measurements.*

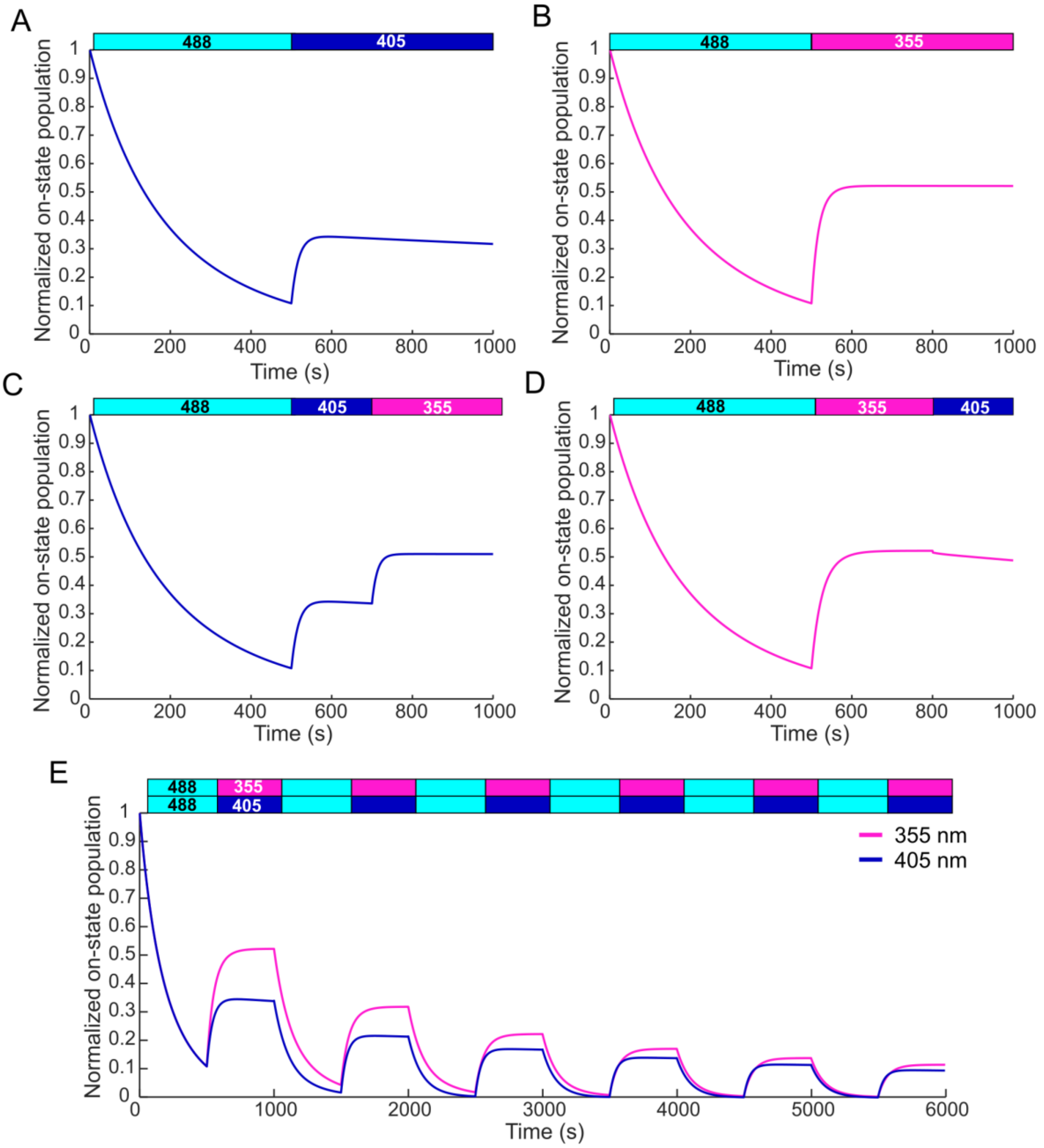


*Figure S16: Ensemble simulations of rsEGFP2 photoswitching at CT. A: Off-switching by 500 W/cm2 488 nm light followed by on-switching by 500 W/cm2 405 nm light. B: Off-switching by 500 W/cm2 488 nm light followed by on-switching by 30 W/cm2 355 nm light. C: Off-switching by 500W/cm2 488 nm light followed by sequential on-switching by 500 W/cm2 405 nm light and 30 W/cm2 355 nm light. D: Off-switching by 500 W/cm2 488 nm light followed by sequential on-switching by 20 W/cm2 355 nm light and 500 W/cm2 405 nm light. E: Photofatigue switching curves under 500 W/cm2 488 nm light and 20 W/cm2 355 nm or 100 W/cm2 405 nm light.*

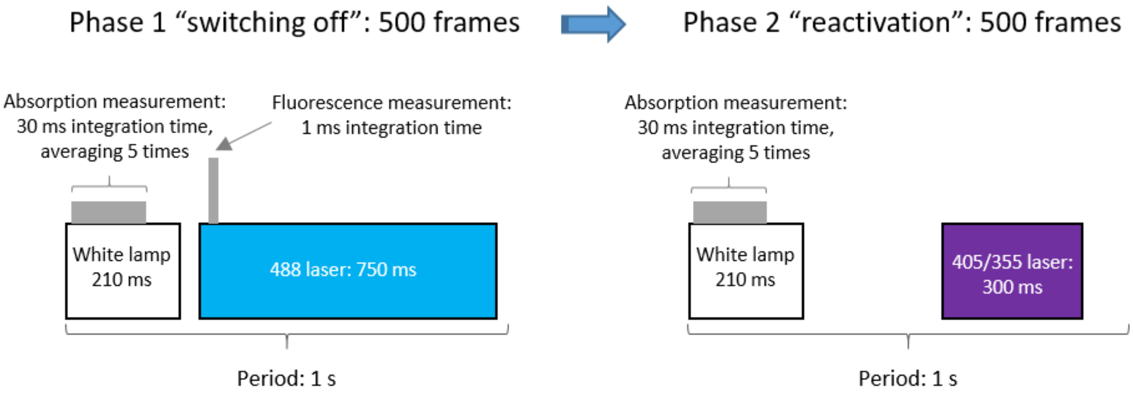


*Figure S17: Illumination scheme employed in microspectrophotometry experiments at CT. During 750 ms of a 1 s period (75% duty cycle) a 488 nm laser was used to switch off the rsEGFP2 molecules, repeated for 500 frames. During the second phase, a 405 nm / 355 nm laser was applied for 300 ms (30% duty cycle), in order to reactivate the molecules, repeated for typically 500 frames. Before laser illumination a white lamp was switched on to measure the absorption of the protein. See methods, section “Solution sample illumination”.*

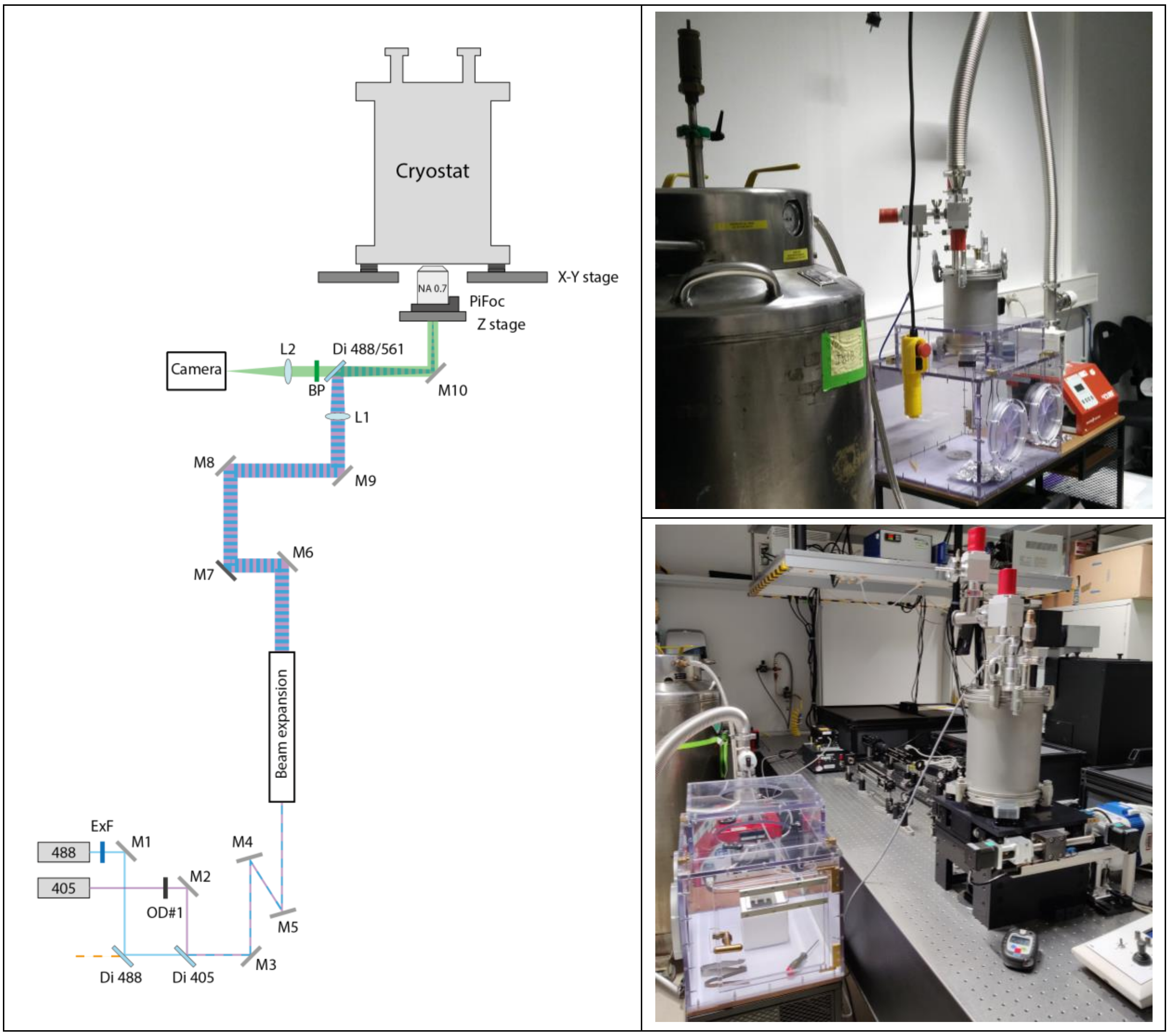


*Figure S28: Schematic illustration and photos of the "cryoPALM" setup. Laser beams at 488 nm (LBX-488-200, Oxxius) and 405 nm (06-MLD 200mW, Cobolt) are first combined together with help of a single-edge 405 nm dichroic mirror (DMLP425, Thorlabs), then expended eight times with a set of achromatic lenses (Thorlabs) and focused with L1 (AC254-300-A-ML, Thorlabs) on the back focal plane of the 60x air objective (LUCPLFLN60X/0.70, Olympus) to produce a wide-field epi illumination of the protein sample mounted in a cryostat. The emission signal is separated from the excitation light with a Di03-R488/561-t1 dichroic mirror (Semrock) and further filtered with a FF01-525/45 band-pass filter (Semrock). The image is finally focused onto an Evolve 512 EM-CCD (Photometrics) camera using an AC508-180-A-ML lens (Thorlabs). A neutral density filter can be inserted into the 405 nm laser beam path for precise adjustment of very low activation laser power. M1 - M10: broadband dielectric mirrors (BB1-E02, Thorlabs).*

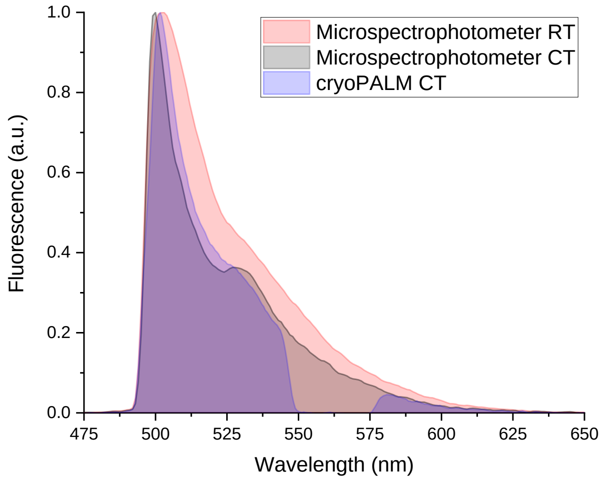


*Figure S19: Fluorescence spectra comparison for measurements done on the micro-spectrophotometer at RT and CT in capillaries and on the cryoPALM setup working with protein-coated cover glasses. The gap in the later one corresponds to the 561 nm reflection band of the Di03-R488/561-t1 dichroic mirror.*

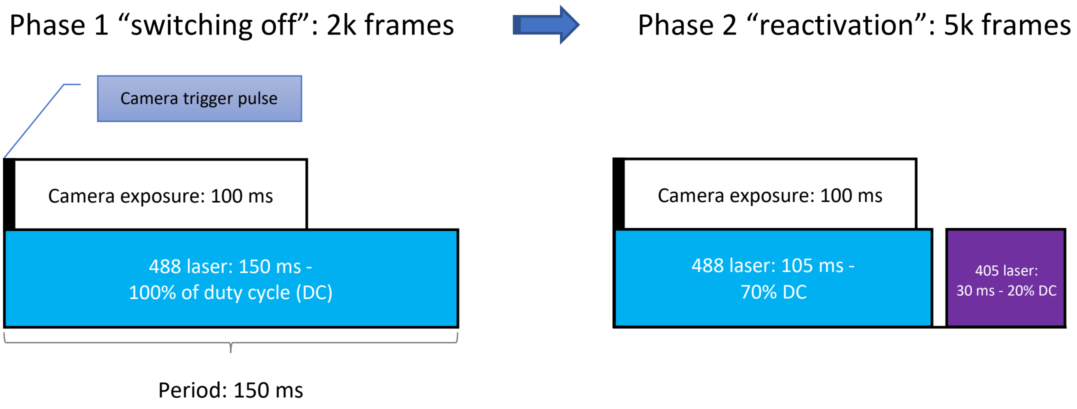


*Figure S20: Illumination scheme employed in single-molecule experiments at CT. During 2000 frames a 488 nm laser was used to switch off the rsEGFP2 molecules and enter the single-molecule regime. During the second phase (5000 frames), a 405 nm laser was added to reactivate the molecules (after the camera exposure to not interfere with image readout).*

### Supplementary Tables

**Table S1. Quantum yields and thermal rates in $s^{-1}$ used for simulations of rsEGFP2 photoswitching at CT.**

| | On1 | On2 | Off1 | Off2 | Offsl_1 | Offsl_2 | Bleached |
|---|---|---|---|---|---|---|---|
| **On1** | - | - | 1.3e-8 | - | 2.6e-5 | - | 1.7e-8 |
| **On2** | - | - | - | 7.1e-8 | - | 2.6e-5 | 7.9e-9 |
| **Off1** | 2.1e-5 | - | - | - | - | - | 1.2e-5 |
| **Off2** | - | 1.7e-6 | - | - | - | - | 1.0e-7 |
| **Offsl_1** | 11.0* | - | - | - | - | - | - |
| **Offsl_2** | - | 11.0* | - | - | - | - | - |

* Thermal rates

Note: Quantum yields and rates are estimations rather than exact calculations.

**Table S2. Data collection and refinement statistics of Crystal 1 (off-switching at CT).**

| Crystal 1 | *Position 1, Initial state* | *Position 2, Cryo-switched off* |
|---|---|---|
| **Pre-illumination** | None | 488 nm, 110K |
| **PDB accession code** | Not deposited | 8AHA |
| **Beamline** | ESRF ID30A-3 (MASSIF-3) | ESRF ID30A-3 (MASSIF-3) |
| **Wavelength (Å)** | 0.9677 | 0.9677 |
| **Resolution range (Å)** | 42.35-2.32 (2.403-2.32) | 46.32-2.38 (2.465-2.38) |
| **Space group** | P 21 21 21 | P 21 21 21 |
| **Unit cell dimensions (a, b, c) (Å)** | 52.46, 60.77, 71.75 | 52.40, 60.75, 71.58 |
| **Unit cell angles (α, β, γ) (°)** | 90.0, 90.0, 90.0 | 90.0, 90.0, 90.0 |
| **Total reflections** | 134796 (13901) | 124380 (12739) |
| **Unique reflections** | 10390 (1021) | 9604 (941) |
| **Multiplicity** | 13.0 (13.6) | 13.0 (13.5) |
| **Completeness (%)** | 99.78 (100.00) | 99.88 (100.00) |
| **Mean I/sigma(I)** | 8.44 (2.31) | 11.44 (2.23) |
| **Wilson B-factor (Å²)** | 34.31 | 33.76 |
| **R-merge** | 0.2258 (1.441) | 0.2481 (1.453) |
| **R-meas** | 0.2352 (1.498) | 0.2583 (1.51) |
| **R-pim** | 0.0649 (0.4044) | 0.07113 (0.4082) |
| **CC1/2** | 0.993 (0.754) | 0.996 (0.714) |
| **CC*** | 0.998 (0.927) | 0.999 (0.913) |
| **Reflections used in refinement** | 10381 (1022) | 9600 (941) |
| **Reflections used for R-free** | 508 (46) | 490 (43) |
| **R-work** | 0.1943 (0.2272) | 0.1857 (0.1997) |
| **R-free** | 0.2436 (0.3260) | 0.2251 (0.3601) |
| **CC(work)** | 0.937 (0.880) | 0.951 (0.886) |
| **CC(free)** | 0.929 (0.694) | 0.918 (0.599) |
| **Number of non-hydrogen atoms** | 2008 | 2008 |
| **macromolecules** | 1886 | 1886 |
| **ligands** | 60 | 60 |
| **solvent** | 62 | 62 |
| **Protein residues** | 237 | 237 |
| **RMS(bonds)** | 0.008 | 0.008 |
| **RMS(angles)** | 1.04 | 1.01 |
| **Ramachandran favored (%)** | 97.00 | 97.85 |
| **Ramachandran allowed (%)** | 3.00 | 2.15 |
| **Ramachandran outliers (%)** | 0.00 | 0.00 |
| **Rotamer outliers (%)** | 0.00 | 0.48 |

| | | |
|---|---|---|
| **Clashscore** | 3.69 | 2.11 |
| **Average B-factor (Å²)** | 36.80 | 36.21 |
| **macromolecules** | 35.89 | 35.29 |
| **ligands** | 65.03 | 65.59 |
| **solvent** | 36.88 | 35.51 |

Statistics for the highest-resolution shell are shown in parentheses.

**Table S3. Data collection and refinement statistics of Crystal 2 (back-switching at CT after off-switching at RT).**

| **Crystal 2** | ***Position 1, RT-switched off*** | ***Position 2, RT-switched off + cryo-switched on*** |
|---|---|---|
| **Pre-illumination** | 488 nm, RT | 488 nm, RT + 405 nm, 110K |
| **PDB accession code** | Not deposited | 8AHB |
| **Beamline** | ESRF ID30B | ESRF ID30B |
| **Wavelength (Å)** | 0.9763 | 0. 9763 |
| **Resolution range (Å)** | 46.40-1.82 (1.885-1.82) | 46.44-1.79 (1.858-1.79) |
| **Space group** | P 21 21 21 | P 21 21 21 |
| **Unit cell dimensions (a, b, c) (Å)** | 51.43, 61.78, 70.26 | 51.67, 61.76, 70.44 |
| **Unit cell angles (α, β, γ) (°)** | 90.0, 90.0, 90.0 | 90.0, 90.0, 90.0 |
| **Total reflections** | 228965 (10899) | 254080 (14673) |
| **Unique reflections** | 20581 (1724) | 21607 (2020) |
| **Multiplicity** | 11.1 (5.7) | 11.8 (7.3) |
| **Completeness (%)** | 98.41 (85.14) | 99.36 (94.43) |
| **Mean I/sigma(I)** | 9.59 (0.31) | 10.87 (0.45) |
| **Wilson B-factor** | 36.04 | 34.59 |
| **R-merge** | 0.1465 (3.939) | 0.1268 (3.356) |
| **R-meas** | 0.1534 (4.348) | 0.1326 (3.616) |
| **R-pim** | 0.04481 (1.785) | 0.03823 (1.315) |
| **CC1/2** | 0.998 (0.155) | 0.999 (0.237) |
| **CC*** | 1 (0.519) | 1 (0.619) |
| **Reflections used in refinement** | 20356 (1724) | 21589 (2018) |
| **Reflections used for R-free** | 1018 (86) | 1081 (101) |
| **R-work** | 0.1959 (0.5422) | 0.1921 (0.4730) |
| **R-free** | 0.2418 (0.5371) | 0.2195 (0.4800) |
| **CC(work)** | 0.962 (0.374) | 0.964 (0.606) |
| **CC(free)** | 0.961 (0.503) | 0.959 (0.555) |
| **Non-hydrogen atoms** | 2030 | 2030 |
| **macromolecules** | 1902 | 1902 |
| **ligands** | 40 | 40 |
| **solvent** | 88 | 88 |
| **Protein residues** | 235 | 235 |
| **RMS(bonds)** | 0.006 | 0.006 |
| **RMS(angles)** | 0.95 | 0.92 |
| **Ramachandran favored (%)** | 99.13 | 99.13 |
| **Ramachandran allowed (%)** | 0.87 | 0.87 |
| **Ramachandran outliers (%)** | 0.00 | 0.00 |
| **Rotamer outliers (%)** | 1.90 | 1.90 |
| **Clashscore** | 3.67 | 2.62 |
| **Average B-factor** | 39.61 | 39.37 |
| **macromolecules** | 38.92 | 38.66 |
| **ligands** | 63.26 | 63.66 |
| **solvent** | 43.69 | 43.61 |

Statistics for the highest-resolution shell are shown in parentheses.
